# Clonal and Scalable Endothelial Progenitor Cell Lines from Human Pluripotent Stem Cells

**DOI:** 10.1101/2023.08.19.553753

**Authors:** Jieun Lee, Hal Sternberg, Paola A. Bignone, James Murai, Nafees N. Malik, Michael D. West, Dana Larocca

**Affiliations:** AgeX Therapeutics Inc., Alameda, CA 94501; Advanced Cell Technology, Alameda, CA 94502; Reverse Bioengineering, Inc., Alameda, CA 94502

**Keywords:** Endothelial progenitor cells, cardiovascular diseases

## Abstract

Human pluripotent stem cells (hPSCs) can be used as a renewable source of endothelial cells for treating cardiovascular disease and other ischemic conditions. Here, we present the derivation and characterization of a panel of distinct clonal embryonic endothelial progenitor cell (eEPC) lines that were differentiated from human embryonic stem cells (hESCs). The hESC line, ESI-017, was first partially differentiated to produce candidate cultures from which eEPC were cloned. Endothelial cell identity was assessed by transcriptomic analysis, cell surface marker expression, immunocytochemical marker analysis, and functional analysis using a vascular network forming assay. The transcriptome of the eEPC lines was compared to various adult endothelial lines as well as various non-endothelial cells including both adult and embryonic origins. This resulted in a variety of distinct cell lines with functional properties of endothelial cells and strong transcriptomic similarity to adult endothelial primary cell lines. The eEPC lines, however, were distinguished from adult endothelium by a novel pattern of embryonic gene expression. We demonstrated scalability of up to 80 population doublings and stable with long-term expansion over 50 passages and stable angiogenic properties at late passage in the EPC line. Taken together, these data support the finding that hESC-derived clonal eEPC lines are useful as a source of scalable therapeutic cells and cell products for treating cardiovascular disease. These eEPC lines offer a highly promising resource for preclinical studies and therapeutic interventions.

## 1. Introduction

Cardiovascular diseases including atherosclerosis, ischemic stroke, myocardial infarction, and ischemic cardiomyopathy currently represent a large economic and societal burden due to their high mortality and disability rates [1]. Impairment of blood vessel formation and maintenance is one of the major underlying causes of many age-associated diseases such as diabetes, cardiovascular disease, peripheral artery disease, and slow wound healing [2]. Aging of angiogenic progenitor populations in adult tissues may account for the lack of homeostatic repair and maintenance of vascular tissues seen with advanced age [3]. Regenerative cell replacement therapies offer a potentially effective strategy for treating conditions involving dysfunctional endothelium. Adult stem cell therapies using mesenchymal stem cells (MSCs) [4, 5], endothelial progenitor cells (EPCs) [6], and cardiosphere-derived cells (CDCs) [7] are currently being investigated as potential therapeutic agents for ischemic diseases [8]. However, the potential of adult stem cell therapies is challenged by issues of identity, scalability, stability, and purity. Human pluripotent stem cells (hPSCs) have the potential to provide a scalable source of nearly all human somatic cell types [9]. Multiple groups have explored the therapeutic efficacy of hPSC-derived endothelial cells (hPSC-ECs) in animal models of ischemic cardiovascular diseases and demonstrated a potential for enhancing angiogenesis, tissue perfusion, and organ graft [10–12]. However, current reports of vascular cell differentiation from hPSC show results in inefficiency (1-5%) [13, 14], heterogeneous aggregates [15] or lack of consistent yields of endothelial cells [16]. Manufacturing hPSC-derived cells at scale presents a problem because of the difficulty reproducing the differentiation protocol for each batch and issues with cost-effective scaling of hPSCs [17]. Current hPSC cell-based therapies in clinical testing such as those being investigated for treating macular degeneration (RPE cells) and spinal cord injury (Oligodendrocytes) require relatively small doses or patient populations [18]. In contrast, cardiovascular disease treatments may necessitate larger doses, and hence more stringent release criteria. Given the heterogeneous nature of hPSC-ECs and their various differentiation protocols, clinical application in disease treatment remains challenging [19, 20].

We have previously developed a method of generating purified and embryonic site-specific somatic cell types through the propagation of hPSC-derived clonal human embryonic progenitor cell lines to overcome current issues of purity and scale [21]. We designated these cultures as human “embryonic progenitor” (hEP) cells because of their ability to self-renew under selected culture conditions, their persistent expression of embryonic developmental stage gene markers such as *PCDHB2*, and their lack of fetal/adult gene markers such as *COX7A1* that are preferentially expressed in cells that have traversed the embryonic-fetal transition [22]. These hEP cell lines also typically display limited lineage potential based on the loss pluripotency markers and pluripotent functionality. In our initial characterization of approximately 200 hEP lines, we reported that they were often capable of robust expansion and displayed a diversity of > 140-fold distinct cell types [21]. Due to the diversity and clonal nature of hEP cell lines, the cells show site-specific markers such as homeobox genes that facilitate the identification of the lines as precursors to specific embryonic anlagen. For example, we characterized adipocyte progenitor cells [21, 23], seven distinct osteochondral progenitor cell types [24, 25], and progenitors of cranial neural crest [26] from our library of hEP cell lines. These hEP cell lines derived from hPSCs were partially differentiated and selected for scalable clones. More importantly, they were able to further differentiate into (1) brown adipocyte cells for metabolic disease treatment, (2) chondrocytes for cartilage regeneration, and (3) cellular components of the choroid plexus, all of which have unique progenitor properties for cell-based therapy [21–23].

In present study, we derived novel hEP cell lines with endothelial properties and characterized a panel of 14 embryonic endothelial progenitor cell (eEPC) lines to explore whether eEPCs can be potential source of therapeutic endothelial cells. We compared the eEPC lines to a diverse panel of adult endothelial cell (AEC) lines using comprehensive transcriptomic analysis and found that we 13 out of 14 eEPC lines expressed a broad array of endothelial specific genes shared across various endothelial subtypes. We assessed eEPC lines for endothelial identity using cell surface markers and well-established vascular tube forming assays. The eEPCs were clearly distinguished from AECs based on embryonic specific gene expression and molecular network analysis. In addition, we were able to achieve high scalability typical of previously identified hEP lines, suggesting that they may be advantageous for developmental research and for treating cardiovascular disease in a large patient population.

## 2. Materials and Methods

### Derivation of Endothelial Progenitor Cell Lines

Research grade hESCs (ESI-017) were differentiated using a two-step progenitor derivation protocol. In the first step, human embryonic stem cells are exposed to a 7-day procedure on Matrigel coated plates involving specific differentiation factors, followed by exposure to two different growth medium to generate two cultures of heterogenous cells designated as Candidate Cultures (CC). After, each CC subpopulation was placed at clonal densities in their same growth medium for two weeks. Then individual clonal colonies were selected with cloning cylinders and expanded from a 24-well plate to multiple T225 flasks (**Figure 1A**).

**Figure 1.**
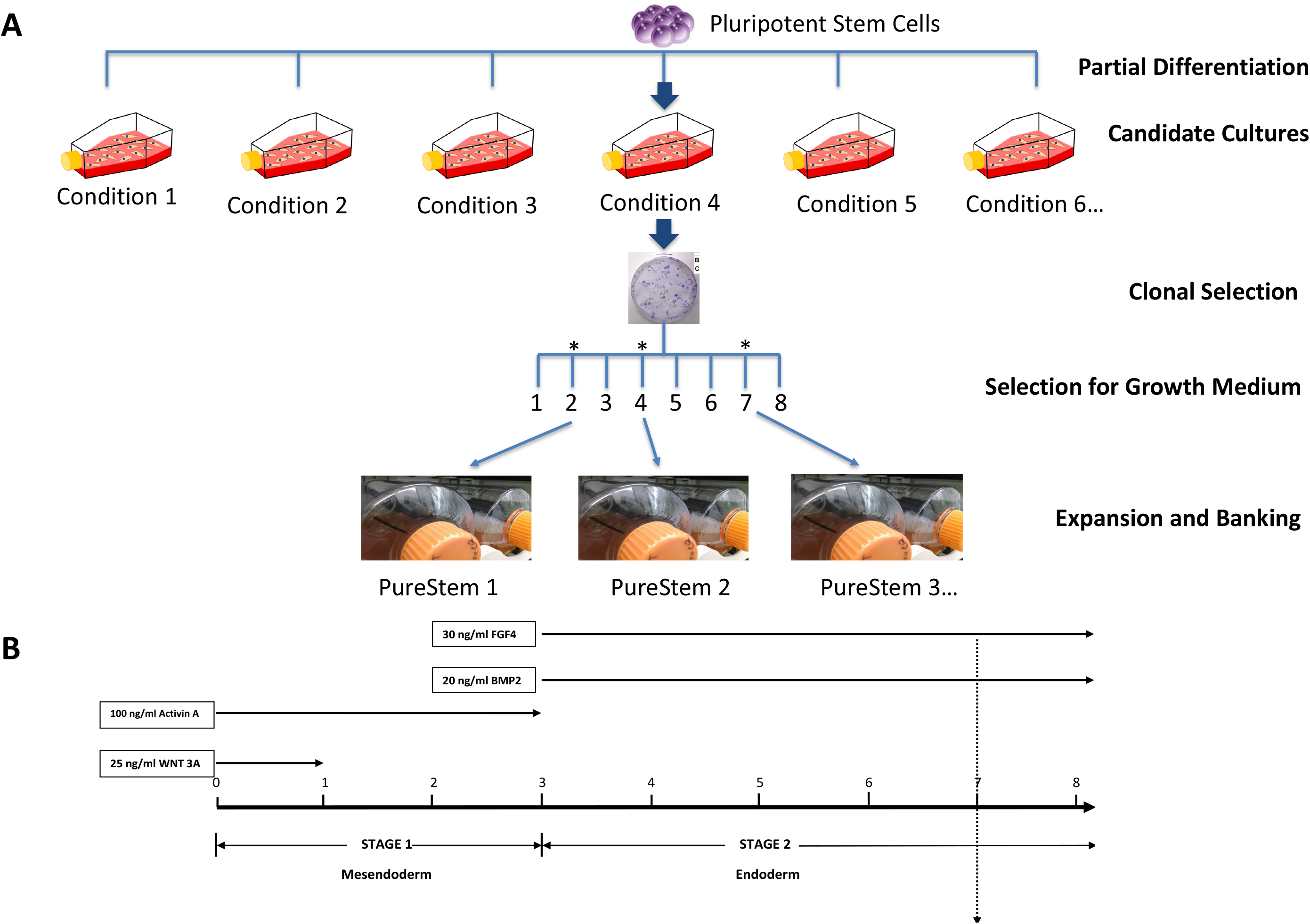
Derivation of Endothelial Progenitor Cell Lines, (A) Schematic illustration of establishing embryonic endothelial progenitor lines from clonal selection and (B) eEPC differentiation strategy from hPSCs, Protocol “30”.

As described above and in **Figure 1** [13], hESCs were differentiated to promote mesoendoderm/endoderm commitment in the presence of specific growth factors for specific period of times to obtain the desired endothelial progenitor characteristics. Briefly, after reaching confluence, hESCs were exposed to: (day 0), basal differentiation medium supplemented with 25ng/ml WNT3A (removed at day 1); and 10 ng/ml activin A (removed at day 3); on day 3, medium was supplemented with 30 ng/ml FGF-2 and 20 ng/ml BMP4 (remained for the duration of culture). On day 7, the differentiated cells were divided into two growth medium as: (1) Microvascular endothelial growth medium (EGM-MV2, Cat# C-39226, PromoCell, GmbH, Germany) with added (TGF) β-inhibitory molecule SB431542 (10 μM) and (2) Smooth Muscle Growth Medium (SM2) (PromoCell, Cat # C22062) with added SB431542 (10uM). After further expansion in the two growth medium, the cells were seeded at clonal dilution for two weeks. Individual clonal colonies were then selected and placed in a 24-well plate (each clonal colony/well given an independent name) and scaled to multiple T225 flasks before they were frozen down and later thawed for further analysis as described in West et al. [21] (**Figure 1B** and **Table 1**). Throughout the expansion and analysis of eEPCs, cells were continually maintained in the presence of SB431542 in their original growth medium that they were exposed after the 7 days differentiation procedure.

**Table 1.** Culture medium content and list of growth factors.

| Basal Differentiation Medium |  |  |  |  |  |
| --- | --- | --- | --- | --- | --- |
| Component | Source | Catalog Number | Lot | Volume | Final Concentration |
| Knock-Out DMEM/F12 | Invitrogen | 10829-018 | 896672 | 500 ml |  |
| Glutamax-1 | Invitrogen | 35050-061 | 930000 | 5 ml | 2 mM% |
| NEAA | Invitrogen | 11140-050 | 815047 | 5 ml | 1% |
| Pen – Strep | Invitrogen | 15070-063 | 932245 | 5 ml | 1% |
| b-Mercaptoethanol | Invitrogen | 21985-023 | 762405 | 0.6 ml | 90 uM |
| RPMI 1640 | Invitrogen | 21870-070 | 1000391 | 100 ml |  |
| FBS | Invitrogen | 26140-079 | 788065 | 200 ul | 10% |
| Stemline II | Sigma | S0192-500ml | 081M442 |  |  |
| Growth Factors |  |  |  |  |  |
| Factor | Source | Catalog Number | Final Concentration |  |  |
| WNT3A | StemRD | W3A-H-005 | 25 ul/ml |  |  |
| Activin A | Humanzyme | HZ-1138 | 100 ng/ml |  |  |
| FGF-4 | Millipore | GF-098 | 30 ng/ml |  |  |
| BMP-2 | Humanzyme | HZ-1129 | 20 ng/ml |  |  |
| SB431542 | Millipore | 616464 | 10 uM |  |  |

### Cell Lines and Culture

14 eEPC lines and 15 human adult endothelial cell (AEC) lines were cultured in endothelial growth medium (EGM-MV2, Cat# C-39226, Lot# 25 x 461M089, PromoCell, GmbH, Germany) on gelatin-coated plates. All the information on cell lines including cell origins and sources were described in **Supplementary Table 1.** The medium was changed every 2-3 days and cells are passaged at 80% confluency. The cells when maintained in the undifferentiated state were cultured at 37°C in a humidified atmosphere of 10% CO2 and 5% O2

### Transcriptomic RNA-sequencing Analysis

RNA was prepared upon lysis with RLT with 1% 2-bME, using Qiagen RNeasy mini kits (Cat#74104) following manufacturer’s directions. The extracted RNA was then quantitated using a NanoDrop (ND-1000) spectrophotometer. Library Construction was performed by using Illumina Truseq mRNA library prep kit following manufacturer’s directions. Library QC and library pooling was accomplished using Agilent Technologies 2100 Bioanalyzer to assay the library fragments. qPCR was used to quantify the libraries. Libraries were pooled, which had different barcodes/indexing and sequencing, in one lane. The paired-end sequencing was performed using the Illumina HiSeq4000 sequencing instrument, yielding 100-bp paired-end reads. The sequencing was performed by BGI Americas Corporation. Data analysis of the transcription levels (FPKM values) was performed using Gene Spring Suite. Hierarchical and correlation analysis was performed with moderate T-test and p (Corr) cut-off=0.05 while applying Pearson metric. The dendrogram was created using hierarchical cluster analysis using average agglomeration method. For each cell culture sample, the differential expression values were calculated against a selection of 14 eEPC clones versus 15 adult endothelial cell lines versus 17 non-endothelial cell lines including embryonic and adult origin. All the information on samples is described in **Supplementary Table 2**. Gene ontology analysis was performed using PANTHER Test from GO Consortium with FDR correction. Raw data used for the current study are available from the corresponding author upon request.

### Whole Genome Microarray Analysis

Total DNA was extracted from cells using Qiagen DNeasy mini kits according to instructions supplied by the manufacturer. DNA concentrations were measured using a Nanodrop spectrophotometer. Whole-genome expression was obtained using Illumina Human HT-12 v4 BeadArrays. In preparation for Illumina BeadArrays, total DNA was linearly amplified and biotin-labeled using Illumina TotalPrep kits (Life Technologies, Temecula, CA, USA). The sample quality was measured using an Agilent 2100 Bioanalyzer before being hybridized to Illumina BeadChips, processed, and read by an iScan microarray scanner according to the manufacturer’s instructions (Illumina, San Diego, CA, USA). Values under 130 relative fluorescence units (RFUs) were considered as nonspecific background signal. Analysis of microarray data was performed using GeneSpring suite. Raw microarray data were normalized with the R beadarray library [27]. Merging of data from different experiments and their subsequent quantile normalization was performed using functions combine and lumiN, respectively, of lumi library. Dendrograms were created by hierarchical cluster analysis, correlation and 3-diemnsional PCA analysis were created in GeneSpring suite.

### Statistical analyses

Statistical significance of data was determined by applying a two-tailed student’s t test to values obtained from independent experiments.

### Flow Cytometry

Cells were dissociated into a single-cell suspension by using TrypLE (Invitrogen-Life Technologies) and fixed in BD Cytofix buffer (BD Biosciences) for 20 min at room temperature. The cells were permeabilized by washing and incubating them with BD Permeabilization/Wash (BD Biosciences) buffer at 1×10^6^ cells per 1 ml for 10 min. The cells were stained by incubating them with antibodies (mouse anti-human Sox9, anti-human CD31-Pacific Blue, and anti-human VE-Cadherin-APC; BDBiosciences) for 30 min. Primary antibodies were diluted according to the manufacturer’s instructions. The cells were scanned with an LSRII flow cytometer (BD Biosciences) and analyzed with FlowJo software (Ashland, OR, USA).

### Immunocytochemistry

For detection of CXCR4 and CD31, cells were washed once with PBS and fixed in 4% paraformaldehyde for 30–60 min at room temperature (RT). Fixed cells were washed three times with PBS, permeabilized and blocked by incubation in blocking buffer (5% normal donkey serum, 1% BSA and 0.1% Triton X-100 in PBS) for 1 h at RT. The cells were then incubated overnight at 4 °C with primary rabbit anti-human CXCR4 polyclonal antibody (Thermo Sci. PA1-24894) at a dilution of 1:500 in 5% normal donkey serum, 0.5% BSA and 0.05% Triton X-100 in PBS. Then, the cells were washed four times with PBS plus 0.05% Triton X-100 (PBS-Triton) and incubated for 1 h at RT with Alexa Fluor 568 donkey anti-rabbit IgG antibody (Invitrogen, A10042) at a 1:500 dilution in PBS-Triton. Isotype controls were stained under identical conditions except that total rabbit IgG (Life Technologies, 10500C) was used as primary antibody. Cells were counterstained with DAPI at 0.1 ng/mL for 10 min at RT and imaged on a Nikon Eclipse TE2000-U inverted microscope.

### Vascular Tube Forming Assay

Tube formation angiogenesis assay was performed using the Cell Player Angiogenesis PrimeKit (Essen Bioscience) to assess tube network growth in eEPC lines. eEPCs were labeled using TagRFP control vector consists of the lentiviral backbone vector, pLKO.1-puro, containing a gene encoding red fluorescence protein (RFP), driven by the CMV promoter (Millipore Sigma, Cat#SHC012). RFP labeled eEPCs were plated with monolayer and 10ng/ml of recombinant human VEGF protein (R&D System, Cat# 293-VE-010) was treated to enhance tube formation. Images were taken every 6 hours for 10-14 days using an IncuCyte imager. Tube formation was quantified using the IncuCtye Angiogenesis Analysis Module by dividing the lengths of all tube networks by the image area at each time point (mm/mm^2^). Exosomes for tube formation assay were isolated from eEPC cultured medium using precipitation method following the manufacturer’s instructions (Invitrogen, Cat#4478359). Total exosome particles were counted using nanoparticle tracking analysis (NTA).

## 3. Results

### Derivation of clonal embryonic endothelial progenitor cell (eEPC) lines

We have previously demonstrated that a large number of highly scalable and clonally pure progenitors can be isolated from human pluripotent stem cells using a two-step process of random differentiation followed by clonal selection of scalable cell lines [21]. In our initial clonal derivation, we isolated over 200 clonal cell lines and found by non-negative matrix factorization that 140 of these were distinct cell types [21]. Our original intention was to first generate a high level of progenitor cell diversity using various initial differentiation conditions followed by clonal selection for producing scalable robust cell lines. For deriving endothelial cell progenitors, we first exposed human embryonic stem cells (hESCs) to directed differentiation conditions, followed by expansion in 2 different growth medium, then seeding at clonal density, clonal selection, and further expansion to obtain scalable endothelial lines [13]. We used both endothelial (RP1) [13] and endoderm (protocol “30”) lineage directed differentiation conditions (**Figure 1 and Supplementary Document 1**). The cell morphology suggested that only the endodermal differentiation conditions (protocol 30) resulted in clonal endothelial lines which was later confirmed by transcriptomic analysis (**Figure 2)**. The cell lines established by the RP1 method in contrast resulted in flat elongated cells, as we previously reported [28]. Whole genome expression analysis comparing clones from RP1 differentiation protocol to clones from “30” differentiation protocol revealed a significant difference in gene expression profile. The cell lines from RP1 method clustered with primary pericyte cell lines (**Supple Figure 1**) [28]. In this current study, we purposed to focus on establishing endothelial progenitor cell lines. In Stage 1 of the method “30” differentiation, we designed to induce mesenchymal to endoderm differentiation with Activin A and Wnt3A treatment for 3 days. Afterwards, in Stage 2, FGF-4 and BMP2 to generate large quantities of HPCs. Based on this differentiation system, we harvested the intermediate progenitor cells according to the surface antigen and performed RNA-sequencing analysis. Clonal lines that were scalable in their selective medium were cultured in progressively larger flasks up to roller bottle culture then harvested and banked.

**Figure 2.**
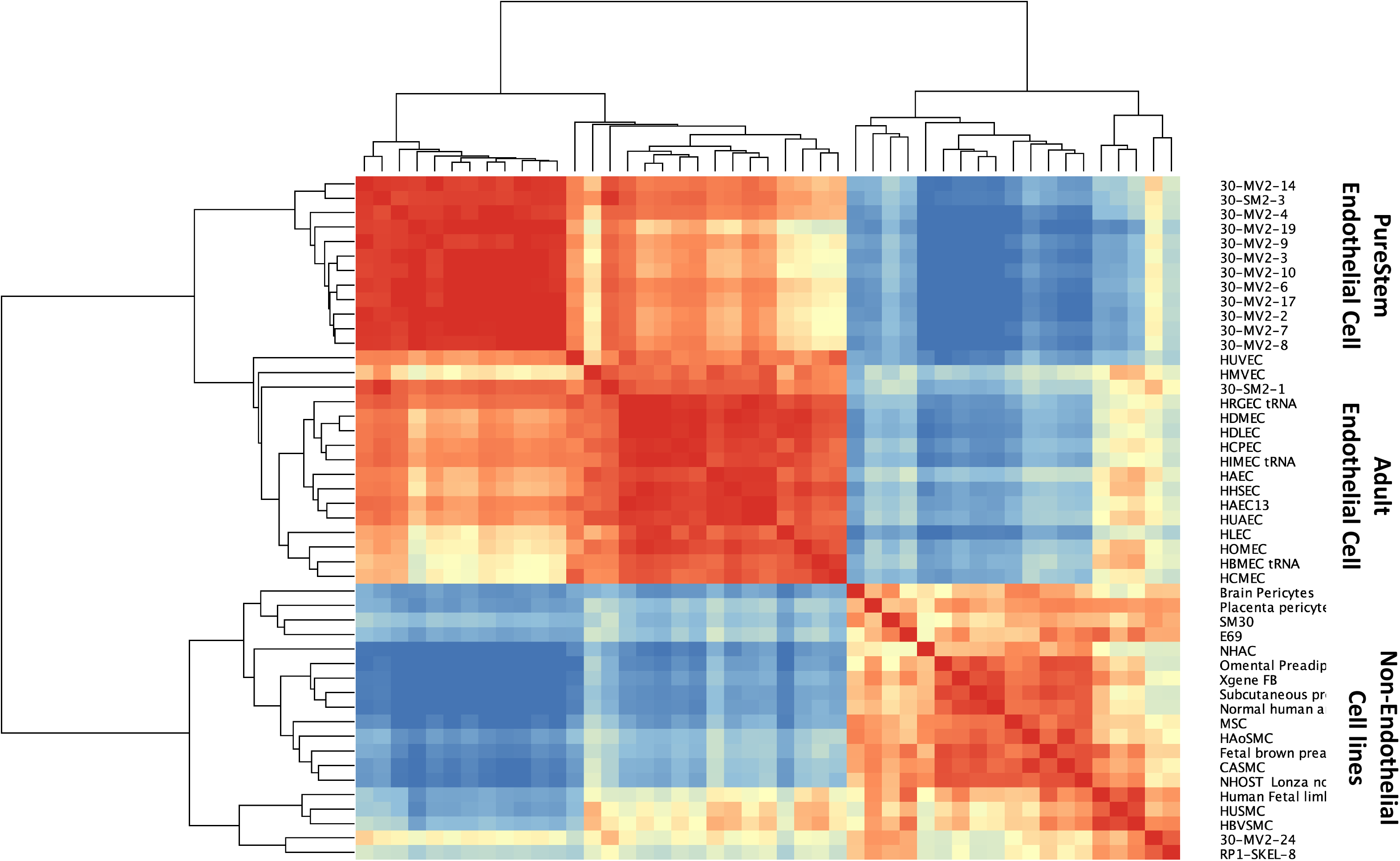
Pearson correlation analysis of (1) human embryonic endothelial progenitor lines (eEPC), (2) adult endothelial cell (AEC) lines and (3) non-endothelial cell lines.

### Transcriptomic analysis indicates the clonal eEPC lines are endothelial cells

To understand the molecular characteristics of the clonally selected hEP lines, we analyzed whole transcriptomic gene expression. We compared three groups: (1) 14 human embryonic endothelial progenitor cell (eEPC) lines, 30-MV2-6, 30-MV2-3, 30-MV2-4,30-MV2-10, 30-MV2-17, 30-MV2-19, 30-MV2-2, 30-MV2-7, 30-MV2-9, 30-MV2-14, 30-MV2-24, 30-MV2-8, 30-SM2-1, and 30-SM2-3; (2) 15 primary adult endothelial cell (AEC) lines including HBMEC, HCPEC, HDMEC, HDLEC, HLEC, HIMEC, HRGEC, HHSEC, HCMEC, HAEC, HOMEC, HUVEC, HUAEC, HMVEC, HAEC, and (3) a panel of 17 non-endothelial cell lines (both embryonic and fetal/adult). The description of cell origin, cell sources, and ages of RNAseq samples were described in **Supplementary Table 1**. Pearson correlation analysis, principal component analysis (PCA), and unsupervised clustering of whole transcriptomic data clearly were performed and the comprehensive analysis indicated that 13 out of 14 eEPC lines clustered with the primary AEC lines, demonstrating a strong correlation between eEPC and AEC lines, whereas both adult and embryonic non-endothelial cell lines were distinct from eEPC and primary AEC lines (**Figure 2** and **Supplementary** Fig.2).

Hierarchical clustering analysis with a representative heatmap in **Figure 3** showed that 13 out of 14 eEPC lines shared a group of endothelial specific genes with primary AEC lines. Pathway enrichment analyses of 399 genes that were commonly expressed in both eEPC and AEC lines was performed. The biological gene ontology (GO) terms showed that endothelial cell differentiation (GO 0045446), vascular development (GO 0048514, GO 0001944 and GO0001568), and angiogenesis (GO 0001525) were significantly enriched, suggesting that 13 clonal eEPC lines express endothelial characteristics comparable to primary endothelial cells.

**Figure 3.**
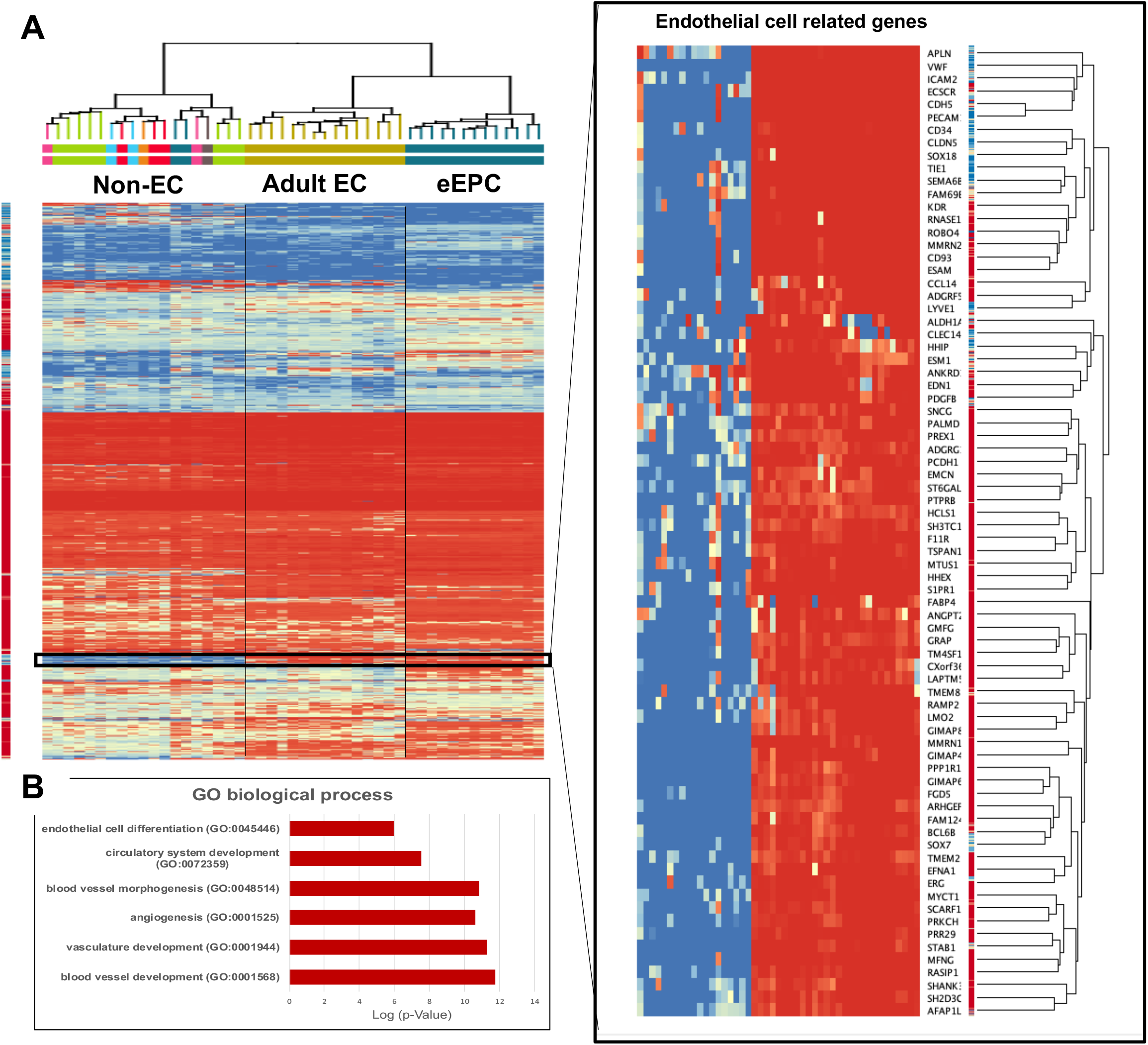
Endothelial characteristics of 14 clonal eEPC lines comparable to 15 human adult endothelial cell lines and 17 non-endothelial cell lines, (A) Transcriptomic RNA-seq hierarchical analysis, moderate T-test and p (Corr) cut-off=0.05, (B) Biological gene ontology (GO)

### eEPC lines express surface proteins and cellular markers of endothelial cells

To further assess the endothelial characteristics of eEPC lines, we initially tested the expression of the key endothelial markers, such as cluster of differentiation (CD) 31, VE-Cadherin (*CD144*), vascular endothelial growth factor receptor (*VEGFR-2)/KDR*, and Von Willebrand factor (*vWF*), which are all widely recognized as endothelial-specific markers [10, 29]. Phase contrast imaging of representative eEPC lines displayed a cobblestone shape morphology (**Figure 4A**) and 99.7% eEPC population expressed *CD31+/ VE-Cadherin +* cells according to single cell flow cytometric analysis (**Figure 4B**). We also confirmed the CD31 expression in a representative eEPC line; 30MV2-6, using immunofluorescence staining (**Figure 4C**) as well as transcriptomic gene expression of endothelial specific genes (*KDR* and *vWF*) (**Supplemental Fig. 3**). Interestingly, we observed that the eEPC lines expressed platelet-derived growth factor-B (PDGF-B), inhibitor of differentiation 1 (ID1), and C-X-C Motif Chemokine Receptor 4 (CXCR4), with comparable level to AEC lines (**Supplemental Fig. 3**). It has been reported that PDGF-B, ID1 and CXCR 4 play a role in stimulating vascular circulation [30], preserving vascular commitment via TGFβ inhibition [13] and regulating tissue regeneration as well as angiogenesis [31], respectively. These data suggested that the eEPC lines possess global gene expression profiles regulating molecular network of endothelial cells.

**Figure 4.**
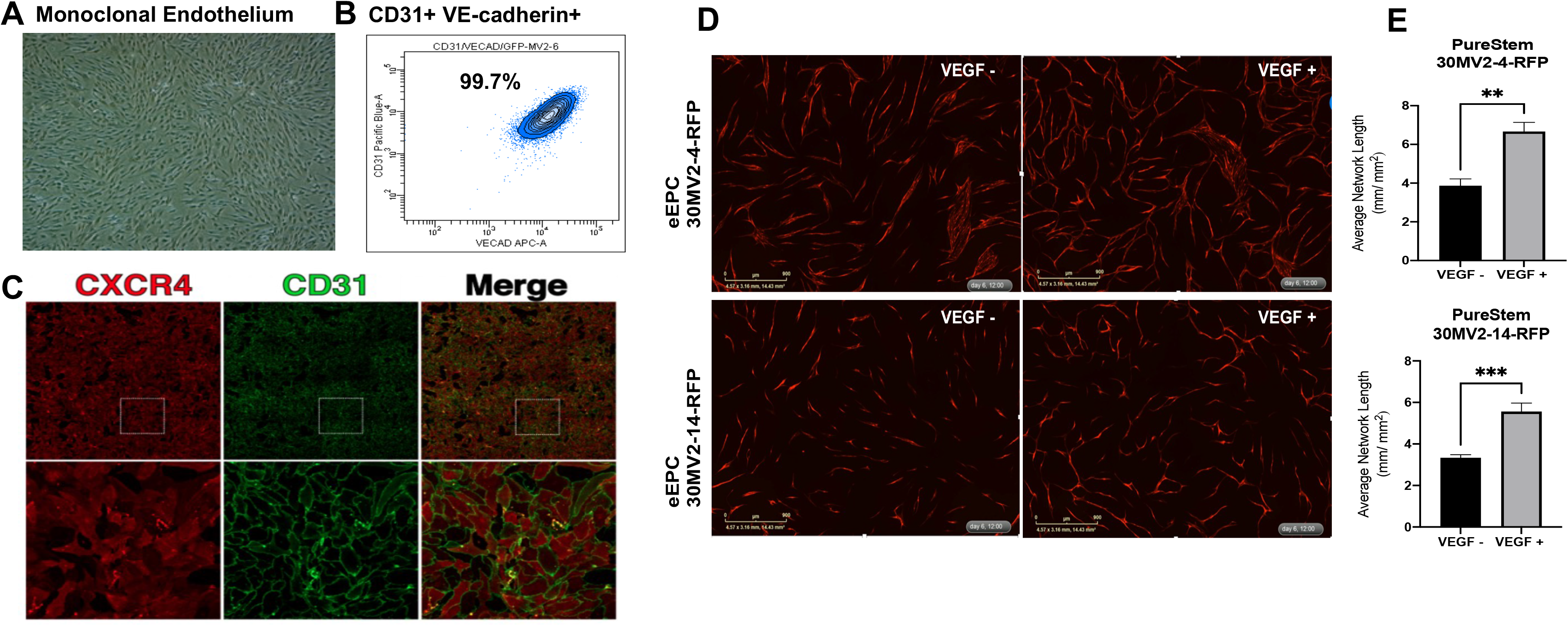
Characterization of clonal endothelial embryonic progenitor cell lines, (A) Phase contrast image of the representative eEPC line, 30MV2-6, (B) Flow cytometric analysis of CD31 and VE-Cadherin, (C) Immunofluorescence staining of endothelial markers, CXCR4 and CD31, and (D) Live cell tube formation angiogenesis assay responding VEGF and quantitative analysis of tube network length of two representative eEPC lines, 30MV2-14 and 30MV2-14.

### eEPC lines enhance tube formation in response to VEGF

We next investigated the angiogenic functionality of eEPC lines for vascular tube formation *in vitro*. We previously demonstrated these eEPC lines differentiated from hEPs were not tumorigenic [21]. To assess tube network growth in eEPC lines, eEPCs were labeled usin*g TagRFP* and co-cultured with normal human dermal fibroblasts (NHDF). This model allows us to demonstrate all phases of the angiogenesis process, including proliferation, migration, and, eventually, differentiation and angiogenic networks [32]. Imaging the co-culture in live-cell analysis system enabled us to identify RFP-labeled eEPCs from co-cultured NHDF and to visualize the vessel formation networks of eEPC over 6 days (**Figure 4D and Supple Video 1**). To quantify the amount of tube formation, we combined time lapse image acquisition to measure network tube length. **Figure 4E** shows tube formation that is responsive to VEGF, an integral proangiogenic cytokine, in two representative eEPC lines (30MV2-4 and 30MV2-14) over 6 days. The angiogenic potency and response to VEGF further supports the endothelial nature of these eEPC lines.

### eEPC lines retain embryonic phenotype

To better understand the molecular signature of eEPC lines compared to AEC lines, we performed a differential gene expression analysis of the transcriptomic RNAseq data. Although the overall global gene expression patterns of the 13 eEPC lines were similar to 15 AEC lines from various origins, we found that there was a significant subset of genes that were differentially expressed in the eEPC lines (**Figure 5**). The RNAseq data were validated by the independent measurement of showing global gene expression profile using microarray analysis (**Supple Figure 4**). To define a specific identity of eEPC lines, differentially expressed genes in eEPCs were compared to genes expressed in AECs and ranked according to the average fold change in transcript expression. Indeed, we observed 1092 genes with at least 2-fold increased expression and 579 genes with at least 2-fold decreased expression in eEPC lines relative to AEC (**Figure 6A**). Pathway enrichment analysis showed that eEPC-enriched genes were related to vascular cell migration, vascular transport, and vessel development (**Figure 6B)**. In contrast, AEC-enriched gene pathways related to regulation of mitochondrial ATP synthesis, mitotic spindle elongation, and metabolic processes. We found a set of genes reported as major endothelial regulators that were highly upregulated in eEPC lines, including Apelin receptor (*APLNR*, differentiation maker of hematopoietic stem and progenitor cells) [33], insulin-like growth factor 2 (*IGF2*, promoting EPC homing) [34], *MafB* (controlling endothelial sprouting) [35], Insulin-like growth factor binding protein 5 (*IGFBP5*, promotes angiogenesis) [36], *H19* (a long non-coding RNA increasing EC tube formation) [37], Carnitine palmitoyltransferase 1 C (*CPT1C*, promoting human mesenchymal stem cells survival) [38], Tartrate-resistant acid phosphatase 5 (*ACP5*, promoting cell differentiation and proliferation) [39], and *EGLN3* (enhanced prolyl hydroxylase domain (PHD)-3, regulator associated with resistance to stress) [40] (**Figure 7A** and **Supple Fig. 5A**).

**Figure 5.**
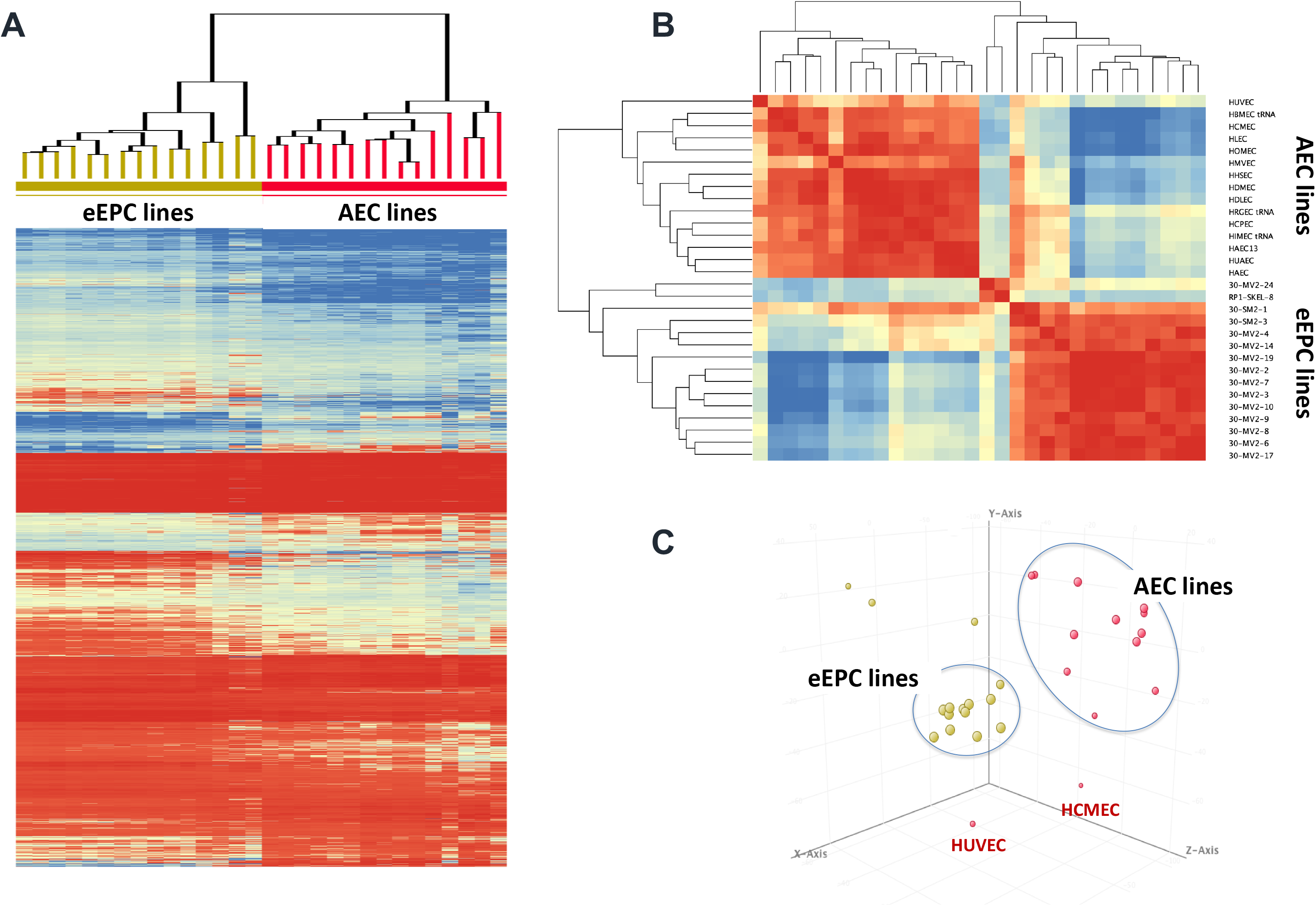
RNAseq Transcriptomic analysis showing retention of embryonic phenotype in eEPC comparing to adult endothelial cell lines. (A) RNAseq cluster analysis heatmap using moderate T-test, (B) Gene Correlation analysis, p (Corr) cut-off=0.05 and (C) principal component analysis (PCA)

**Figure 6.**
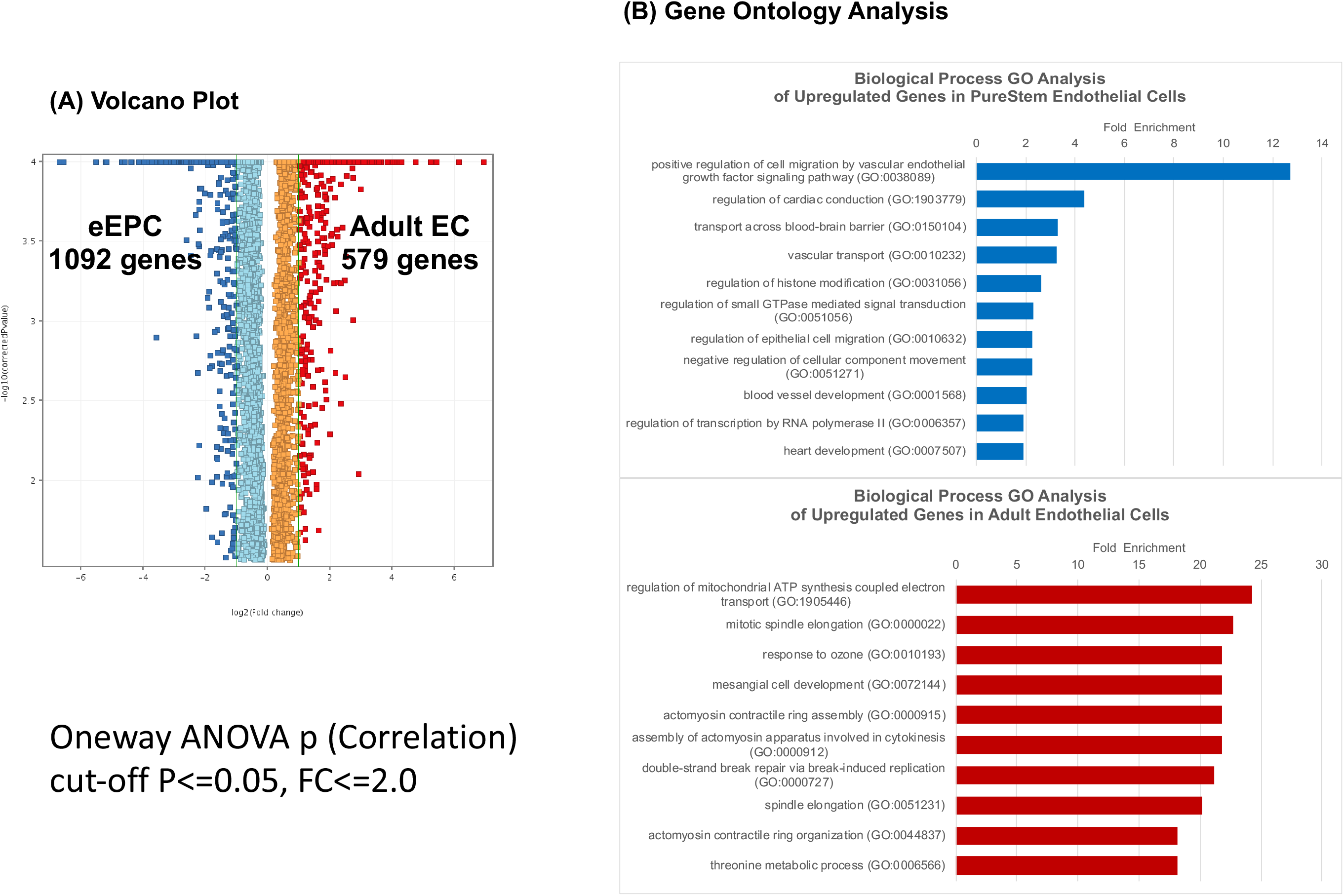
Differential gene expression analysis of Adult EC lines vs. eEPC lines. (A) Volcano Plot using One-way ANOVA p (Correlation), cut-off P<=0.05, Fold Channge<=2.0 and (B) Gene Ontology Analysis

**Figure 7.**
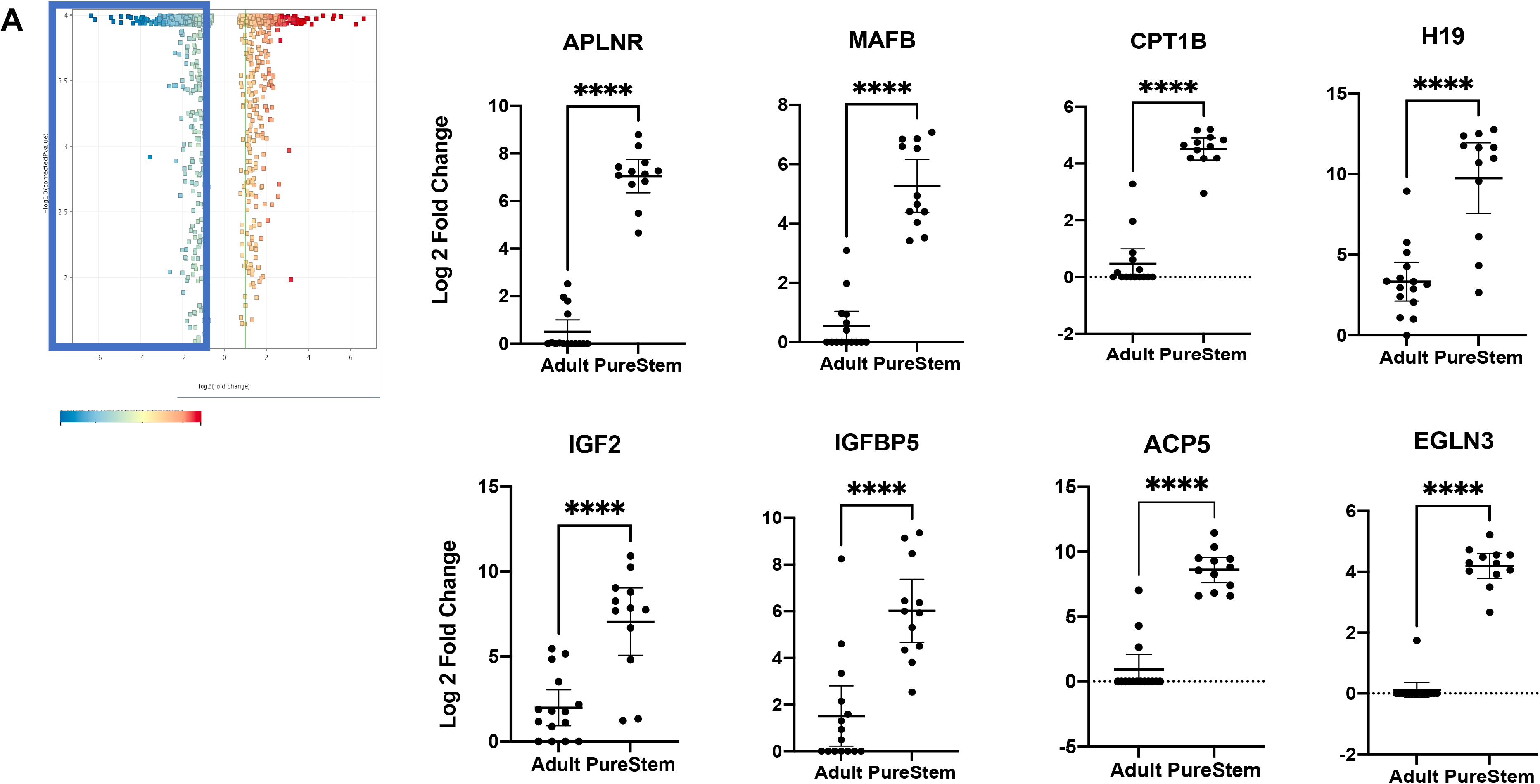
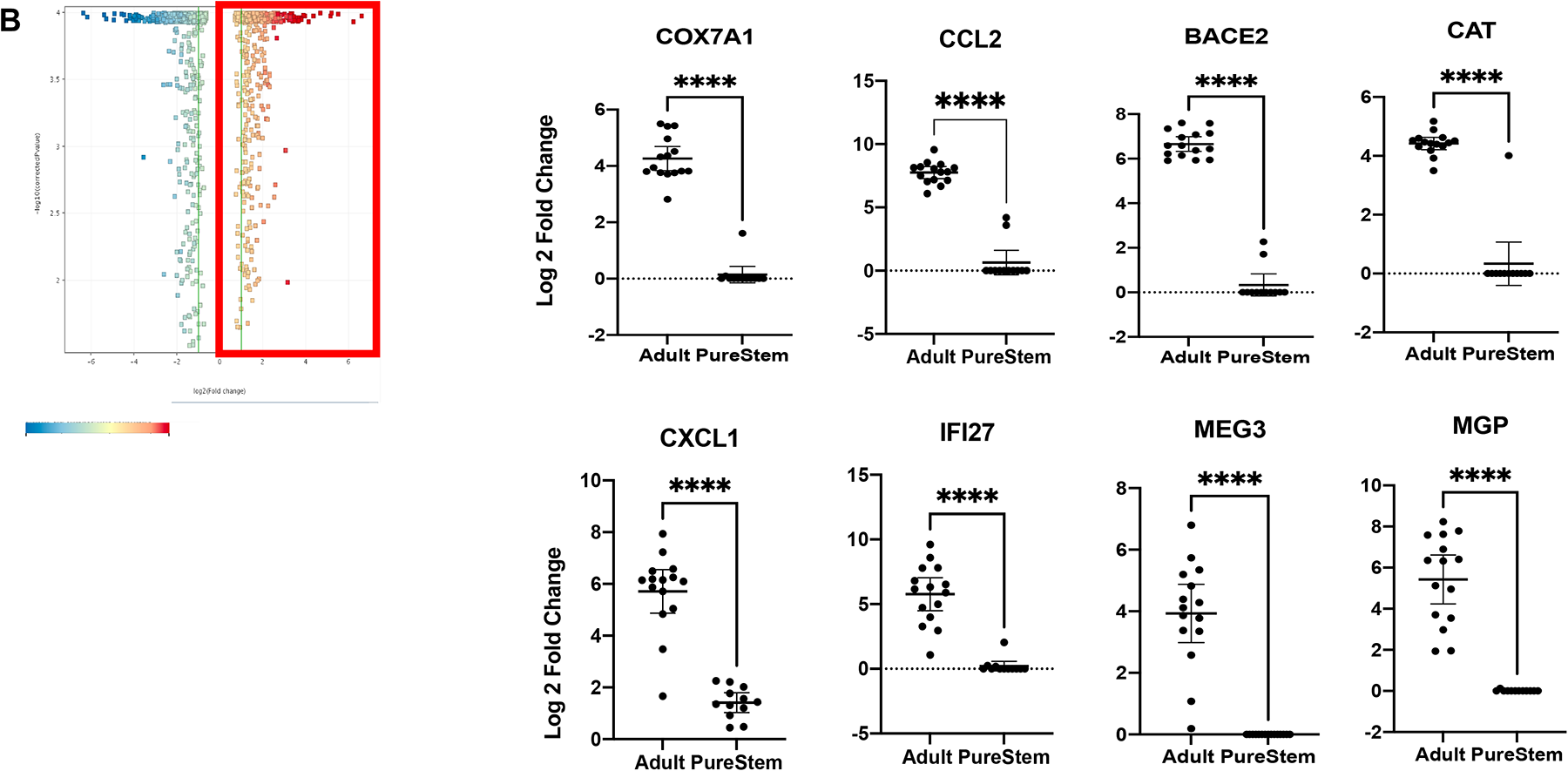
List of genes of. (A) Highly expressing genes expression in eEPC lines comparing to Adult EC lines and (B) Highly expressing genes expression in Adult EC lines comparing to eEPC lines, Oneway ANOVA p (Correlation), cut-off P<=0.05, FC<=2.0

We previously identified *COX7A1*, encoding a cytochrome C oxidase subunit, as a novel marker associated with the mammalian embryonic-fetal transition (EFT) that is almost exclusively expressed in fetal and adult cells [22]. *COX7A1* expression level in eEPC lines was observed to be undetectable (Log2 FC= 0.1), compared to AEC lines (Log2 FC= 4.3), suggesting that eEPC lines retained embryonic progenitor stage (**Figure 7B**). In addition, the gene expression of CC chemokine ligand 2 (*CCL2*, signaling associates with diabetes) [41], β-site APP cleavage enzyme 2 (*BACE2*, gene that are linked to increased risk and earlier disease onset such as Alzheimer’s disease) [42], *CAT* (an antioxidant enzyme, play an important role in endothelial cell shear stress response) [43], chemokine (C-X-C motif) ligand 1 (*CXCL1*, regulated in TNF-stimulated endothelial cells) [44], Interferon α-inducible protein 27 (*IFI27*, a stimulator of VEGF-A mediated angiogenesis) [45], Matrix Gla protein (*MGP*, regulates differentiation of endothelial cells) [46], and *MEG3* (long non-coding RNA regulating angiogenesis) [47] were AEC specific and not detectable in eEPC lines (**Figure 7B** and **Supple Fig. 5B**). Taken together, we identified a subset of genes that were differentially expressed in eEPCs compared to AECs, suggesting that eEPCs, while clearly expressing an endothelial gene network, have not undergone terminal endothelial differentiation to fetal/adult but instead express an embryonic pattern of gene expression.

### Stable production of angiogenic embryonic endothelial progenitor cells (eEPCs)

We examined cell growth characteristics by measuring scalability and functional stability of a representative eEPC line, 30-MV2-6. It has been suggested that EPCs have therapeutic potential and clinical implications in ischemic diseases [6, 48–50]. Therefore, the stable and scalable production of EPCs can be beneficial for supporting preclinical and clinical research on eEPC-based therapy.

To test cell production capacity of eEPC lines, cumulative population doublings (PDs) of cells was compared to primary MSCs. We found that MSCs begin to senesce and lose proliferative and differentiation capacity after 12 population doublings (pd) consistent with previous reports [51, 52]. In contrast the eEPC line showed an exponential growth up to at least 80pd (**Figure 8A).** To assess the angiogenic activity of eEPCs, we used eEPC conditioned medium by testing a tube formation assay. The conditioned medium from a representative eEPC line, 30-MV2-6 was precipitated to isolated secreted vesicles from the cells. The treatment of eEPCs-secreted vesicles in human umbilical vein endothelial cells (HUVECs) resulted in enhancing tube formation, which was equivalent to complete endothelial cell growth medium (CGM) containing VEGF, IGF and FGF. These data suggest that extracellular vesicles (also called exosomes) secreted from eEPCs contain angiogenic factors. Recent studies have shown that exosomes are secreted from a variety of cell types including MSCs. They play a role in intercellular communication through the transfer of their cargo carrying lipids, proteins, and RNAs to recipient cells [53]. In this study, we initially observed that exosomes secreted from 30MV2-6 cells stimulated vascular network formation (**Figure 8B)**. The angiogenic activity of 30MV2-6 exosomes was compared to BM-MSC exosomes and tested at a dose of 200 x 10^6^ exosomes/well (**Supple Figure 6A**). The results showed that angiogenic activity was greater for 30MV2-6 than BM-MSC exosomes (P<0.01).

**Figure 8.**
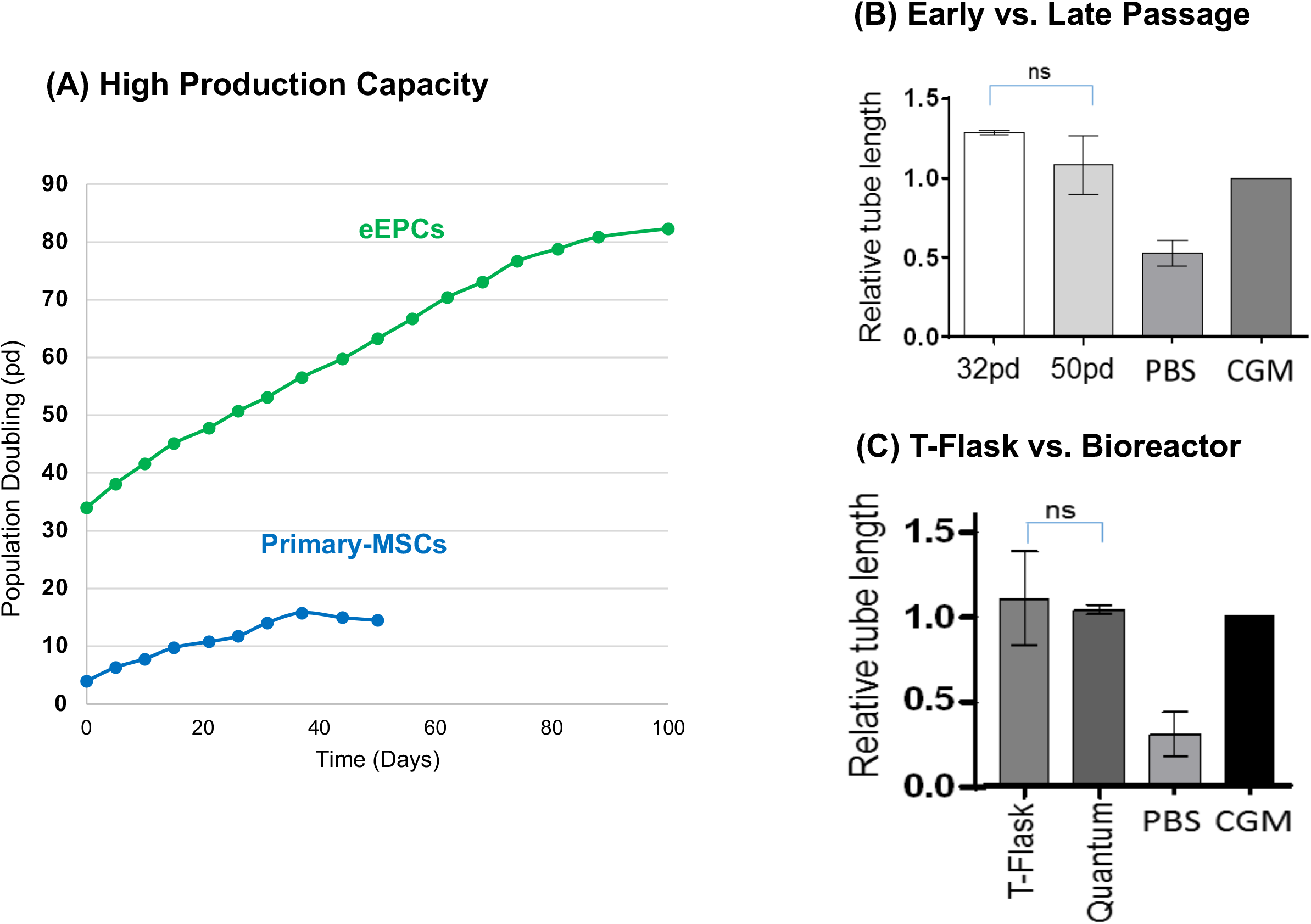
Stable production of embryonic endothelial progenitor cells (eEPCs), (A) eEPC line was scalable up to 80 population doublings comparing to primary MSCs, in contrast, begin to senesce and lose proliferative and differentiation capacity after 8 to 12 population doublings, (B) eEPCs-secreted vesicles enhanced tube formation in vitro and angiogenic activity retained comparing 30pd to 50pd; CGM (EC growth factor cocktails), and (C) Scale-up in a Quantum bioreactor of EPC expansion also resulted in no loss of angiogenic activity.

We next tested whether BM-MSC exosomes were active at a higher dose. A dose of 1200 x 10^6^ exosomes/well of BM-MSC exosomes were equivalent angiogenic activity as 30MV2-6 exosomes at 200 x 10^6^ exosomes/well indicating a 6-fold difference in potency (**Supple Fig. 6B**). These data demonstrated the potential for highly scalable eEPCs to produce exosomes with higher angiogenic potency than BM-MSC. Furthermore, we identified miR-126 to be the single most highly enriched miRNA in 30MV2-6 compared to BM-MSC exosomes by qPCR showing a 50-fold enrichment of 30MV2-6 (**Supple Fig. 6C**), confirming the angiogenic potential of eEPC secreted exosomes. miR-126 is well established for proangiogenic activity and promotes maturation to functional vessels [53–55]. These data suggest an important role for enriched miR-126 in the higher angiogenic activity of eEPC-exosomes and in their potential to improve functional outcomes in the treatment of ischemic disease.

We also compared eEPCs-exosomes cultured from early (32 pd) versus late passages (50 pd). The results show that the early passage eEPC exosome angiogenic activity was stable at later passage (**Figure 8B)**. To investigate the scalability of eEPCs, we compared the angiogenic activity of eEPCs grown in monolayer culture to Quantum bioreactor (Terumo) culture after incubation in endothelial basal medium (EBM) for 72 hours at 5% oxygen. The large-scale eEPC culture in the cell expansion bioreactor had equivalent angiogenic tube formation activity as conventional monolayer culture (**Figure 8C**). Taken together, we demonstrated stability and scalability of eEPCs, showing that large-scale expansion and late passage eEPCs maintained their angiogenic activity. This finding suggested that eEPC lines may be able to overcome major obstacle of primary cells on the path to producing endothelial cells at industrial scale for therapeutic applications.

## 4. Discussion

Previous studies have suggested that mesenchymal stem cells (MSCs) [4, 5], endothelial progenitor cells (EPCs) [6], and cardiosphere derived cells (CDCs) [7] are attractive cell sources for vascular regeneration. However, their widespread use has been limited by their heterogeneity and lack of scalability. Endothelial cells differentiated from hPSCs (hPSC-EC) can be generated by overcoming the major bottleneck of the paucity of cells and donor heterogeneity, but these technologies have also met limitations for therapeutic usage due to heterogeneous cell populations [56]. The aim of the current study was to derive homogeneous, clonally scalable, and well-defined human endothelial cell lines. Here, we provide detailed analysis of global gene expression patterns comparing eEPC lines to adult EC (AEC) lines. The eEPC and AEC lines share a pan-endothelial specific gene expression pattern, however, the eEPC lines are distinguished from adult lines by a distinct expression of a subset of embryonic specific genes, indicating that they retain embryonic characteristics. Embryonic progenitor lines which have not crossed the embryo-fetal transition have longer replicative lifespans and may have greater regenerative properties as we have previously proposed [21, 22]. The endothelial stem cell identity of eEPC is also indicated by the expression of hematopoietic stem cells markers (CD34 and CD133) and endothelial cell markers such as CD31, KDR, vWF, Ve-Cadherin/CD144, Tie2, c-kit/CD117, and CD62E (E-selectin) [57, 58] and CD45, CXCR4, CXCR2, and CCR2. Adult EPCs from different sources express different surface markers. For example, bone-marrow derived EPCs (BM-EPCs), peripheral blood-derived EPCs (PB-EPCs), and cord-blood derived EPCs (CB-EPCs) express different markers [59, 60]. We found that each eEPC line has a distinct gene expression pattern indicating that they may represent progenitors of specific endothelial cell types based on anatomical location.

In the present study, we provided comprehensive transcriptomic profiling analysis demonstrating that eEPCs differentiated from hEPs have endothelial phenotypes expressing endothelial markers such as *CD31*, *CXCR4, KDR, and vWR*, as well as generating tube formation *in vitro.* Interestingly, our study showed that eEPC lines retained embryonic progenitor characteristics. We found a subset of genes that were upregulated in eEPC lines compared to AEC lines. For example, eEPC overexpressed apelin receptor (*APLNR*), insulin-like growth factor 2 *(IGF2*), the MAF BZIP transcription factor B (*MAFB*), insulin-like growth factor binding protein 2 (*IGFBP2),* non-coding RNA *H19,* carnitine palmitoyltransferase 1 C *(CPT1C),* tartrate-resistant acid phosphatase 5 (*ACP5) and* enhanced prolyl hydroxylase domain (PHD)-3 (*EGLN3)* relative to AEC. *APLNR* signaling is required for the generation of cells that give rise to HSCs [33] and there is a prominent role of the *IGF2*/*IGF2R* system in promoting EPC homing [34]. The *MAFB* gene encodes a transcription factor that controls endothelial sprouting *in vitro* and *in vivo* [35] and plays an important role in the embryonic development of the lymphatic vascular system [61, 62]. *IGFBP2* is known as a developmentally regulated gene that is highly expressed in embryonic and fetal tissues and markedly decreases after birth [63]. It promotes angiogenic and neurogenic differentiation potential of dental pulp stem cells [36]. *H19* is expressed during EC differentiation and has functional effects on EC tube formation [37]. *CPT1C* promotes human mesenchymal stem cells survival under glucose deprivation through the modulation of autophagy and is strongly associated with the epithelial-mesenchymal program [38]. *ACP5* is upregulated by transforming growth factor-β1 (*TGF-β1*) and subsequently enhances β-catenin signaling in the nucleus, which promotes the differentiation, proliferation, and migration of fibroblasts [39]. EGLN3, a member of the EGLN family of prolyl hydroxylases, increases during skeletal myoblast differentiation [64] and is associated with resistance to stress [40]. Upregulated genes in eEPCs are known to regulate endothelial function, especially during differentiation and cellular stress conditions. Further studies on the biochemical and functional properties of eEPC lines may identify novel cell types for specific therapeutic and diagnostic purposes.

We also found a subset of genes that downregulated in eEPCs. For example, eEPC lines expressed little to no β-site APP cleavage enzyme 2 (*BACE2)*, catalase (*CAT*), CXCL1, Interferon α-inducible protein 27 (*IFI27*), matrix Gla protein (*MGP*), and long non-coding RNA (lncRNA) maternally expressed gene 3 (*MEG3*) compared to AEC lines. The *BACE2* gene is linked to increased risk and earlier disease onset of Alzheimer’s disease. *CAT* encodes an antioxidant enzyme that plays an important role in endothelial cell pathophysiology, in shear stress response, and ultimately, in arterial aging [43]. *CXCL1* was only expressed by old osteoblasts but not young cells [65] and was known to be produced mainly by TNF-stimulated endothelial cells (ECs) as a potent chemoattractant receptor of innate immune system [44]. *IFI27* is known as an oncogene, a strong stimulator of angiogenesis, by increasing secretion of vascular endothelial growth factor (*VEGF-A*) [45]. *MGP* stimulates differentiation of endothelial cells [46], but also has been found in aortic calcified lesions of aging animal model [66]. *MEG3* negatively regulates angiogenesis and proliferation in vascular endothelial cells. *MEG3* overexpression was significantly suppressed by inhibition of miR-9 resulting in proliferation and *in vitro* angiogenesis in vascular endothelial cells [47]. Downregulated genes in eEPCs are associated with cellular organization and metabolic process. The suppression of genes associated with growth inhibition and aging is consistent with embryonic nature of the eEPCs.

Several different sources of endothelial cells have been explored for their ability to revascularize in ischemic injuries, heal wounds, or build microvasculature in engineered tissues. Human umbilical vein endothelial cells (HUVECs) have been a robust source of ECs with proven capability of forming stable microvessels, which are capable of capillary morphogenesis compared to hPSC-ECs [56]. From our comprehensive transcriptomic analysis comparing 13 eEPC lines to 15 AEC lines, we observed that HUVECs expressed a distinct gene expression pattern distinguished from the 14 adult EC lines consistent with their early newborn/fetal nature (**Figure 5** and **Supple Figure 4**). Indeed, a subset of genes were expressed both in HUVEC and the 13 eEPC lines. These include *IFITM1, NNAT, H19, PLVAP,* and *IGF2*, which were highly upregulated and *CYTL1, KRT7, CXCL1,* and *CXCL8,* which were downregulated in HUVECs and eEPC lines, but not the AEC lines (**Supple Figure 7**). *H19* is expressed during EC differentiation and has functional effects on EC tube formation [37]. Our findings comparing eEPC gene expression profiles to HUVEC suggest that HUVECs represent a cell state that is in between embryonic and adult. The increased regenerative capacity of HUVEC compared to AEC is consistent with our *somatic restriction* concept that cells lose regenerative capacity in stages starting with the embryo to fetal transition and continuing through the fetal-newborn and adult transitions [22]. More studies are needed to compare the regenerative capacity of eEPC, HUVEC and AEC lines.

Derivation of eEPC lines that were homogeneous clonal cells with batch-to-batch consistency production provides a unique competitive advantage for unparalleled industrial scale production for therapeutic application in various ischemic diseases. The competitive advantages of eEPC lines are (1) identity and homogeneity, (2) scalability and stability, and (3) diversity (allowing selection of optimal production and high potency.

There are several groups establishing adult (non-embryonic) stem cells such as EPCs, MSCs, and CDCs for clinical application in the treatment of ischemic disease. However, the source of these primary cell lines results in cellular heterogeneity both within a donor line and between donor lines. Thus, there is a manufacturing bottleneck for the consistent production of therapeutic ECs on an industrial scale needed to treat a large cardiovscular disease patient population. We addressed this issue by developing a bank of progenitor cell lines derived by early-stage clonal isolation from partially differentiated hPS cells [21]. This method has resulted in a diverse collection of distinct cell lines with increased homogeneity, stability and scalability because of their clonal purity and long embryonic telomere length [21]. The resulting lines feature homeobox gene expression consistent with an endoderm, ectoderm, mesoderm, or neural crest origin [26]. We have identified a variety of cell fates including bone, cartilage, smooth muscle cells, pericytes, endothelial cells, white and brown adipose tissue, with various applications in regenerative medicine [21, 24–26, 67]. The diversity that we have generated in our hEP line library allows us to select for cell lines that are tailored to specific applications. Interestingly, our endoderm differentiation method led to successful endothelial cell line isolation. An endodermal origin of endothelial cell has recently been reported [68, 69]. We can therefore identify candidate lines based on cell line stability and scalability in addition regenerative potency. Using a directed differentiation variation on our previous method [21, 24–26, 67], we generated 46 embryonic endothelial progenitor cell (eEPC) lines, giving us a unique opportunity to select and develop proprietary stem cell lines for therapeutic application.

## 5. Conclusions

Here, we demonstrated the ability to derive clonally pure endothelial cells from hPS cells (eEPCs) using our 2-step process. We derived 13 eEPC lines and characterized them as endothelial by transcriptomic analysis, biochemical markers, and vascular function. Further, we have shown early evidence of the angiogenic potential of exosomes secreted from eEPC lines. We are currently developing eEPC exosomes manufacturing methods and investigating the potential therapeutic uses of eEPC exosomes for treating ischemic conditions like stroke by improving vascular and neurological recovery. Our findings suggest that eEPC lines will be a valuable source of both cells and exosomes for use in therapeutic vascular regeneration as well as drug screening.

## Supporting information

included in manuscript file

## Author Contributions

For research articles with several authors, a short paragraph specifying their individual contributions must be provided. The following statements should be used “Conceptualization, DL, HS and MW.; methodology, HS, PB, JM.; software, JL.; validation, JL and HS; formal analysis, JL.; investigation, JL and DL.; resources, HS and MW.;; writing and original draft preparation, JL and DL.; writing—review and editing, JL, DL, HS, NM and MW.; visualization, JL.; supervision, MW.; project administration, JL, DL and HS.; funding acquisition, MW. All authors have read and agreed to the published version of the manuscript.” Please turn to the <u>CRediT taxonomy</u> for the term explanation. Authorship must be limited to those who have contributed substantially to the work reported.

## Patents

Exosomes from Clonal Progenitor Cells, 2019 (US 10,240,127); Methods and compositions for targeting progenitor cell lines, 2016 (US Patent 9,175,263)

## Institutional Review Board Statement

NA

## Informed Consent Statement

NA

## Data Availability Statement

Data is available upon request.

## Supplementary Materials

The following supporting information can be downloaded at: www.mdpi.com/xxx/s1,

**Supplementary Table 1.** Information of cell lines for RNAseq including cell origins and sources.

**Supplementary document 1.** RP1 Differentiation Method: Based on Rafii method James et al

**Supplementary Figure 1.** PCA analysis of hEP (human embryonic clonal progenitor) lines (Related to Figure 2)

**Supplementary Figure 2.** RP1 and 30 series microarray data analysis for screening cell lines, (A) PCA, **(A)** (B) Sample Correlation, (C) Hierarchical Clustering, (D) Volcano Plot comparing 30MV2 vs RP1-MV2, (E) Gene Ontology Enrichment

**Supplementary Figure 3.** Endothelial specific gene expression in eEPC lines and adult EC lines (Related to Figure 3)

**Supplementary Figure 4.** Microarray analysis for global gene expression comparison between embryonic EC and adult EC lines, Filtered on Error-CV < 50.0 percent

**Supplementary Figure 5. (A)** Highly expressing genes expression in eEPC lines comparing to AEC lines and (B) Highly expressing genes expression in AEC lines comparing to eEPC lines (Related to Figure 7)

**Supplementary Figure 6.** eEPC exosomes have greater angiogenic potency than BM-MSC exosomes. **(A)** Angiogenic activity of eEPC line, 30-MV2-6 exosomes is higher than BM-MSC and PBS control (p<0.01; ANOVA)). **(B)** Dose response indicates 30-MV2-6 exosomes are 6-fold more potent than BM-MSCs. (n=3, biological replicate experiments in triplicate). **(C)** 50-fold enrichment of miR-126 in 30MV2-6 vs. BM-MSC by qPCR.

**Supplementary Figure 7.** Gene expression profile comparing to HUVEC

**Supple document 1.** RP1 Method: Based on Rafii method James et al

**Supplementary Table 1**. RNAseq Sample list

**Supple Video 1A.** 30MV2-4-RFP tube formation at 20% O2

**Supple Video 1B.** 30MV2-14-RFP tube formation at 20% O2

