## Supplementary material for "Clonal and Scalable Endothelial Progenitor Cell Lines from Human Pluripotent Stem Cells": included in manuscript file

**Supplementary Table 1.** Information of cell lines for RNAseq including cell origins and sources.

| Samples | Origins | Source | Ages |
|---|---|---|---|
| Fetal brown preadipocytes (Zenbio) P4 ctrl RS_29 | Fetal brown preadipocytes | Adipocyte | Fetal |
| Omental Preadipocyte Zenbio (Lot SLOM-18) P9 ctrl RS_31 | Omental Preadipocyte | Adipocyte | Adult |
| Subcutaneous preadipocytes Zenbio (Lot SLOO54) P6 ctrl RS_32 | Subcutaneous | Adipocyte | Adult |
| HBMEC (Human Brain Microvascular Endothelial Cell total RNA), 10 µg RS_1 | HBMEC tRNA | Adult EC | Adult |
| HCPEC (Human Choroid Plexus Endothelial Cell total RNA), 10 µg RS_2 | HCPEC | Adult EC | Adult |
| HDMEC (Human Dermal Microvascular Endothelial Cell total RNA),10 µg RS_3 | HDMEC | Adult EC | Adult |
| HDLEC (Human Dermal Lymphatic Endothelial Cell total RNA), 10 µg RS_4 | HDLEC | Adult EC | Adult |
| HLEC (Human Lymphatic Endothelial Cell total RNA), 10 µg RS_5 | HLEC | Adult EC | Adult |
| HIMEC (Human Intestinal Microvascular Endothelial Cell total RNA), 10 µg RS_6 | HIMEC | Adult EC | Adult |
| HRGEC (Human Renal Glomerular Endothelial Cell total RNA), 10 µg RS_9 | HRGEC | Adult EC | Adult |
| HHSEC (Human Hepatic Sinusoidal Endothelial Cell total RNA), 10 µg RS_11 | HHSEC | Adult EC | Adult |
| HCMEC (Human Cardiac Microvascular Endothelial Cell total RNA), 10 µg RS_12 | HCMEC | Adult EC | Adult |
| HAEC (Human Aortic Endothelial Cell total RNA), 10 µg RS_13 | HAEC13 | Adult EC | Adult |
| HOMEC (Human Ovarian Microvascular Endothelial Cell total RNA), 10 µg RS_15 | HOMEC | Adult EC | Adult |
| HUVEC (Human Umbilical Vein Endothelial Cell total RNA), 10 µg RS_16 | HUVEC | Adult EC | Adult |
| HUAEC (Human Umbilical Artery Endothelial Cell total RNA), 10 µg RS_17 | HUAEC | Adult EC | Adult |
| HMVEC (Human Microvascular Endothelial Cell) P6 RS_19 | HMVEC | Adult EC | Adult |
| HAEC (Human aortic endothelial cells) P6 ctrl RS_38 | HAEC | Adult EC | Adult |
| HUSMC (Human smooth muscle cell) P5 RS_20 | HUSMC | Adult Smooth Muscle | Adult |
| CASMC P14 RS_21 | CASMC | Adult Smooth Muscle | Adult |
| HBVSMC (Human Brain Vascular Smooth Muscle Cells) P3 RS_28 | HBVSMC | Adult Smooth Muscle | Adult |
| HAoSMC (aortic smooth muscle PromoCell) lot 4002012.2 P5 ctrl RS_42 | HAoSMC | Adult Smooth Muscle | Adult |
| NHAC P6 RS_18 | NHAC | Articular Chondrocyte | Adult |
| Human Fetal limb skin FB 8wk (ABR9745) P2 RS_10 | Human Fetal limb skin FB | FB | Adult |
| Xgene FB P17 ctrl RS_33 | Xgene FB | FB | Adult |
| Normal human arm skin fibroblast 82 yr (Coriell GM01706 A) P12 RS_85 | Normal human arm skin | FB | Adult |
| MSC P5 RS_27 | MSC | MSC | Adult |
| E69 P16 ctrl RS_52 | E69 | Neural Crest | Embryonic |
| DPSC P6 ctrl RS_40 | DPSC | None | Adult |
| EN13 P14 ctrl RS_43 | EN13 P14 ctrl RS_43 | None | Embryonic |
| 4D20.8 P15 ctrl RS_44 | 4D20.8 P15 ctrl RS_44 | None | Embryonic |
| 7SMOO32 P11 ctrl RS_46 | 7SMOO32 P11 ctrl RS_46 | None | Embryonic |
| E15 P19 ctrl RS_47 | E15 P19 ctrl RS_47 | None | Embryonic |
| SK11 P16 ctrl RS_48 | SK11 P16 ctrl RS_48 | None | Embryonic |
| 7PEND24 P21 ctrl RS_49 | 7PEND24 P21 ctrl RS_49 | None | Embryonic |
| MEL2 P17 ctrl RS_50 | MEL2 P17 ctrl RS_50 | None | Embryonic |
| T42 P17 ctrl RS_51 | T42 P17 ctrl RS_51 | None | Embryonic |
| C4ELS5.1 P14 ctrl RS_53 | C4ELS5.1 P14 ctrl RS_53 | None | Embryonic |
| E3 P12 ctrl RS_54 | E3 P12 ctrl RS_54 | None | Embryonic |
| ESI004 NP110 SM P12 ctrl RS_55 | ESI004 NP110 SM P12 ctrl | None | Embryonic |
| ESI004 NP88 SM P12 ctrl RS_56 | ESI004 NP88 SM P12 ctrl | None | Embryonic |
| NHOST Lonza normal human osteoblasts) P5 ctrl RS_39 | NHOST Lonza normal | Osteoblast | Adult |
| Brain Pericytes P6 RS_25 | Brain Pericytes | Pericyte | Adult |
| Placenta pericytes P7 RS_26 | Placenta pericytes | Pericyte | Adult |
| 30-MV2-6 P6 RS_70 | 30-MV2-6 | PureStem EPC | Embryonic |
| 30-MV2-3 P6 RS_71 | 30-MV2-3 | PureStem EPC | Embryonic |
| 30-MV2-4 P6 RS_72 | 30-MV2-4 | PureStem EPC | Embryonic |
| 30-MV2-10 P6 RS_73 | 30-MV2-10 | PureStem EPC | Embryonic |
| 30-MV2-17 P6 RS_74 | 30-MV2-17 | PureStem EPC | Embryonic |
| 30-MV2-19 P6 RS_75 | 30-MV2-19 | PureStem EPC | Embryonic |
| 30-MV2-2 P6 RS_76 | 30-MV2-2 | PureStem EPC | Embryonic |
| 30-MV2-7 P6 RS_77 | 30-MV2-7 | PureStem EPC | Embryonic |
| 30-MV2-9 P6 RS_78 | 30-MV2-9 | PureStem EPC | Embryonic |
| 30-MV2-14 P6 RS_79 | 30-MV2-14 | PureStem EPC | Embryonic |
| 30-MV2-24 P6 RS_80 | 30-MV2-24 | PureStem EPC | Embryonic |
| 30-MV2-8 P6 RS_81 | 30-MV2-8 | PureStem EPC | Embryonic |
| 30-SM2-1 P6 RS_82 | 30-SM2-1 | PureStem EPC | Embryonic |
| 30-SM2-3 P6 RS_83 | 30-SM2-3 | PureStem EPC | Embryonic |
| RP1-SKEL-8 P6 RS_84 | RP1-SKEL-8 | PureStem EPC | Embryonic |
| SM30 P15 ctrl RS_45 | SM30 | PureStem SM | Embryonic |

**Supplementary document 1. RP1 Differentiation Method: Based on Rafii method James et al**

1. Day -1: split (with accutase) ESI 017 on matrigel 3x10e6 cells into StemCells Aggrewell (24 well) and spin to make EB aspirate and add fresh Stemline 2, 1ml/well.
2. Day 0: Stemline 2 with BMP4 20ng/ml
3. Day 1: Stemline 2 with final BMP4 20ng/ml, and activin A 10ng/ml
4. Day 2: Stemline 2 with 8ng/ml FGF2, BMP4 20ng/ml, and activin A 10ng/ml .
5. Day 3: Stemline 2 with 8ng/ml FGF2, BMP4 20ng/ml, and activin A 10ng/ml .
6. **Day 4**: Place on **matrigel** with Stemline 2 with 8ng/ml FGF2, VEGF A 25ng/ml and BMP4 20ng/ml.
7. Day 5:  Stemline 2 with 8ng/ml FGF2, VEGF A 25ng/ml, and BMP4 20ng/ml.
8. Day 6: Stemline 2 with 8ng/ml FGF2, VEGF A 25ng/ml, and SB431542 10uM
9. Day 7: Split onto matrigel in various medium with SB431542 10uM in 6-well and scale.

**Reference: Nature Biotechnology 28 (2):161-6.**

**Supplementary Figure 1. RP1 and 30 series microarray data analysis for screening cell lines**

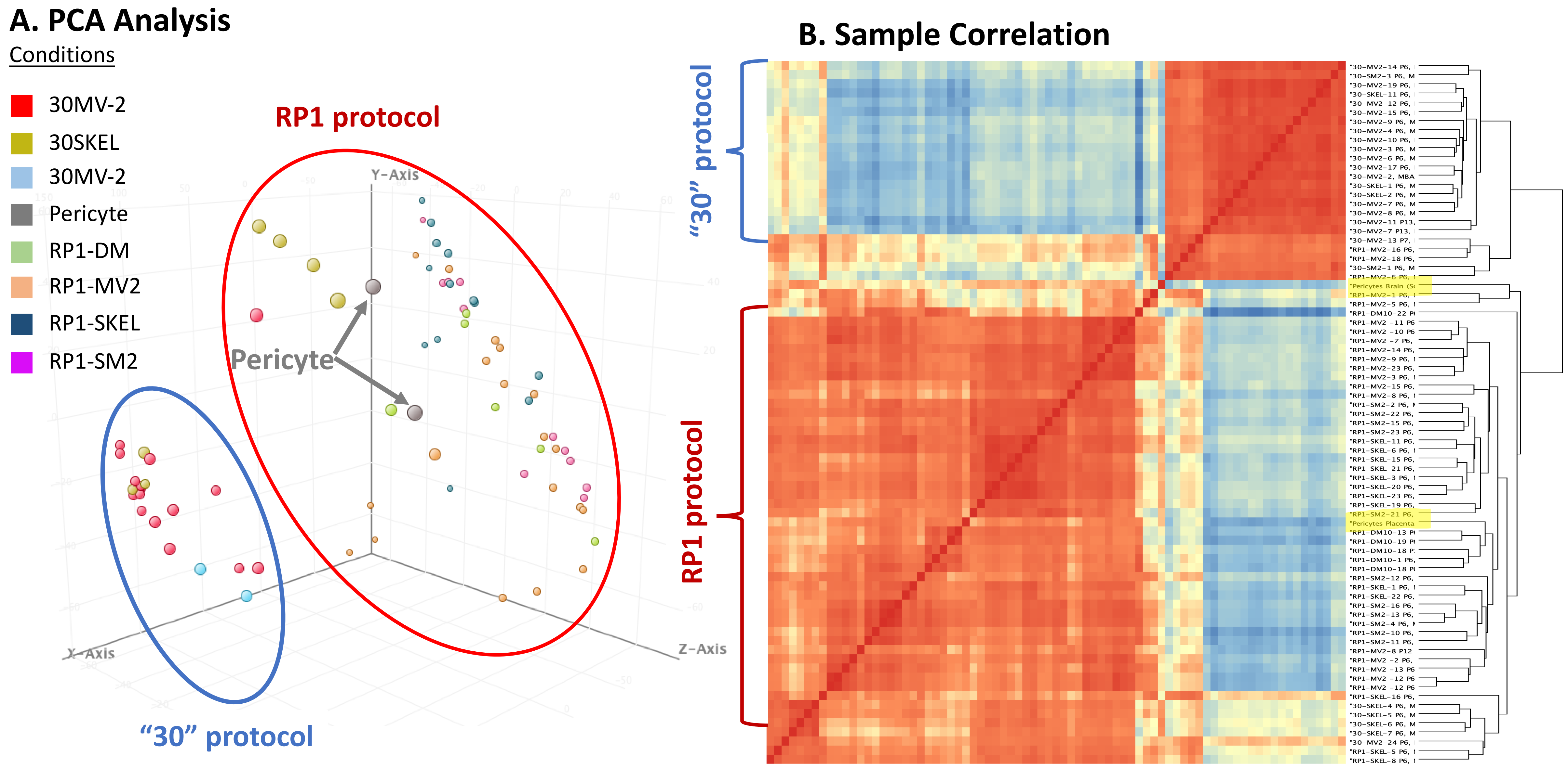

**Supplementary Figure 1. RP1 and 30 series microarray data analysis for screening cell lines**

### C. Hierarchical Clustering

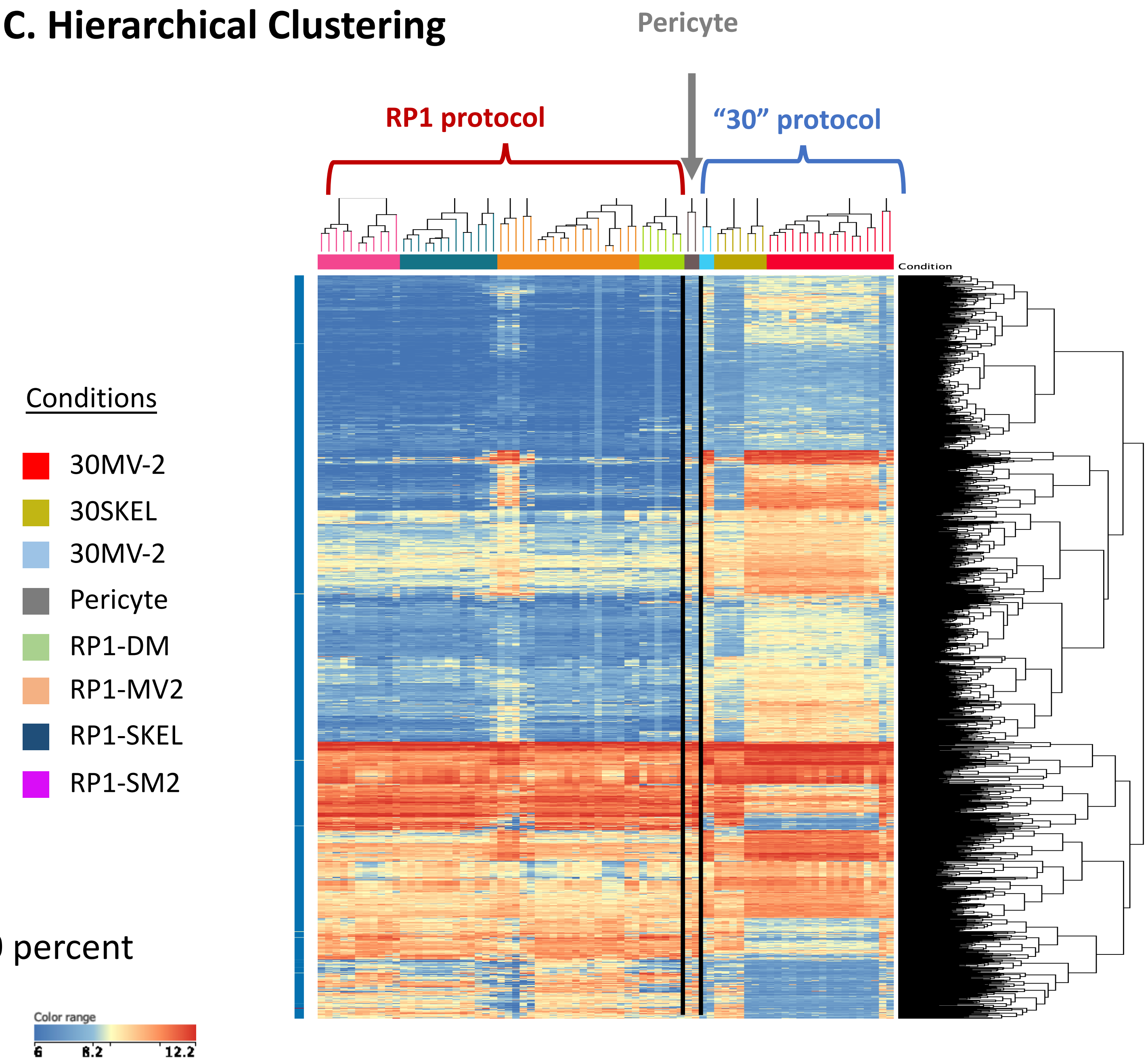


Filtered on Error CV< 50 percent
P<=0.05, FC>=2.0

**Supplementary Figure 1. RP1 and 30 series microarray data analysis for screening cell lines**

**D. Volcano Plot comparing 30MV2 vs RP1-MV2**

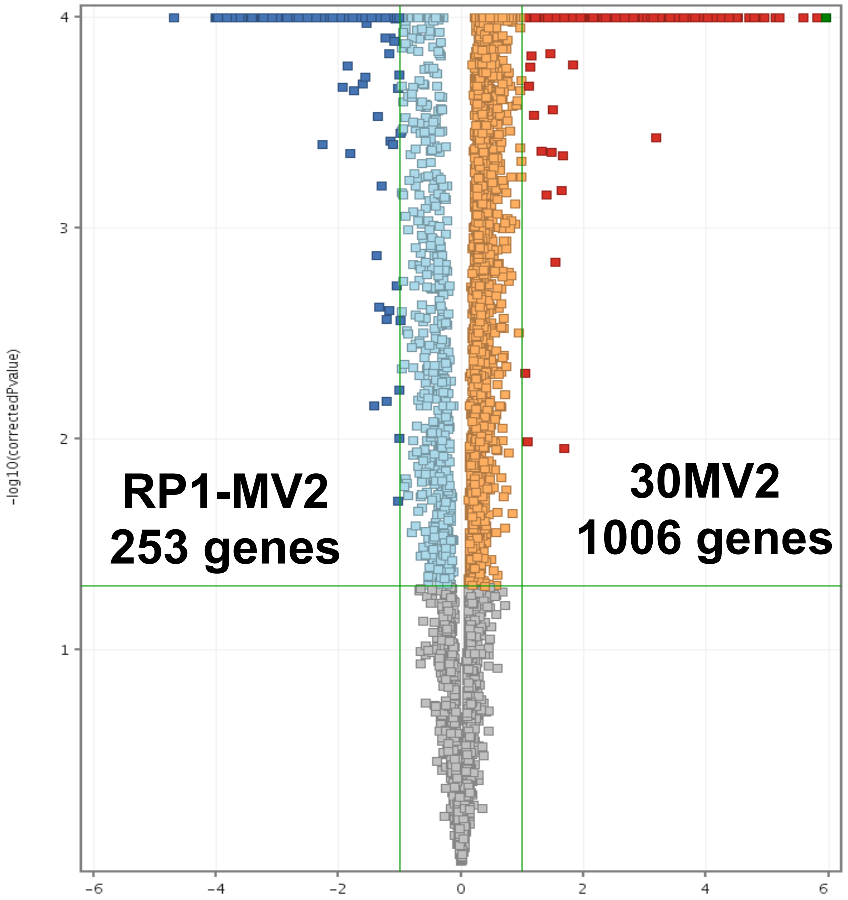


Oneway ANOVA p (Correlation)
cut-off P<=0.05, FC<=2.0

**E. Gene Ontology Enrichment**

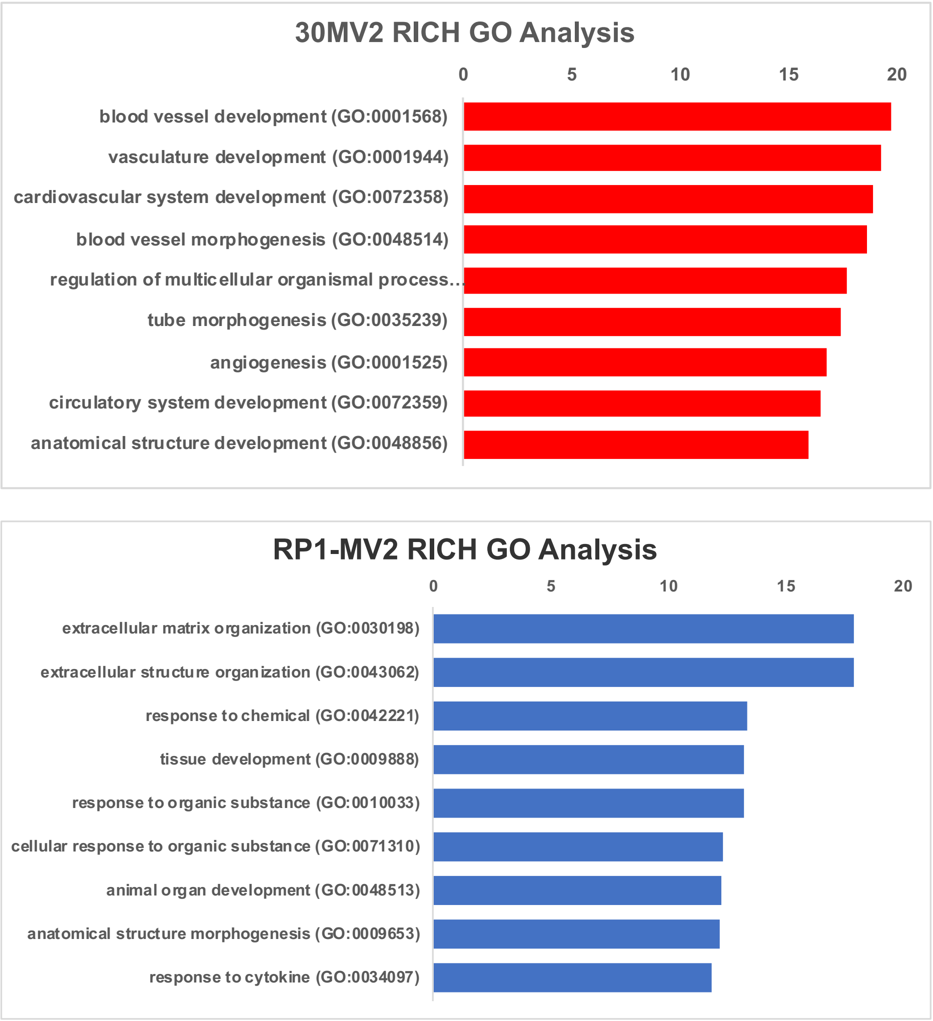


http://geneontology.org/

**Supplementary Figure 2. PCA analysis of hEP (human embryonic clonal progenitor) lines (Related to Figure 2)**

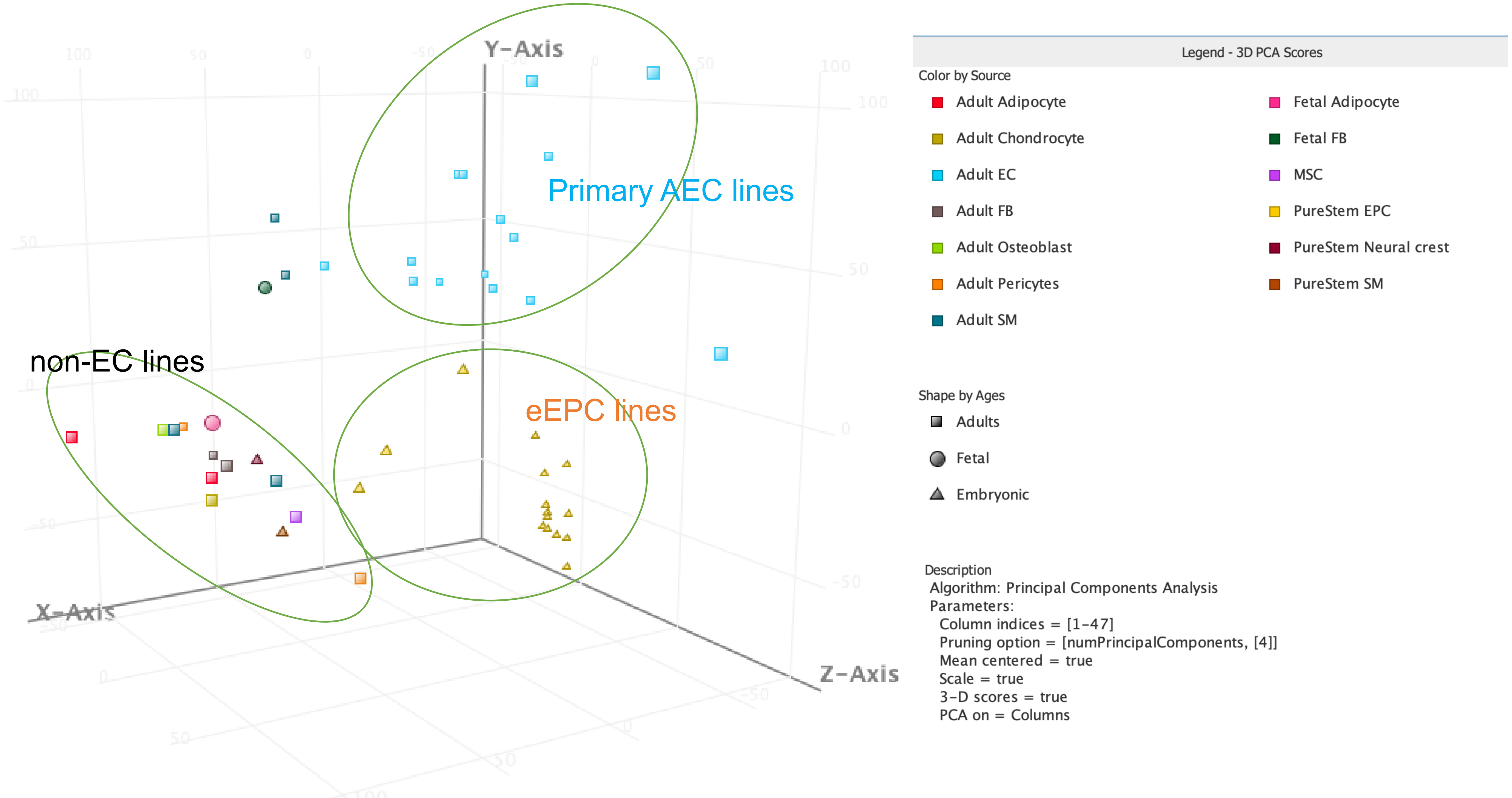

**Supplementary Figure 3. Endothelial specific gene expression in eEPC lines and adult EC lines (Related to Figure 3)**

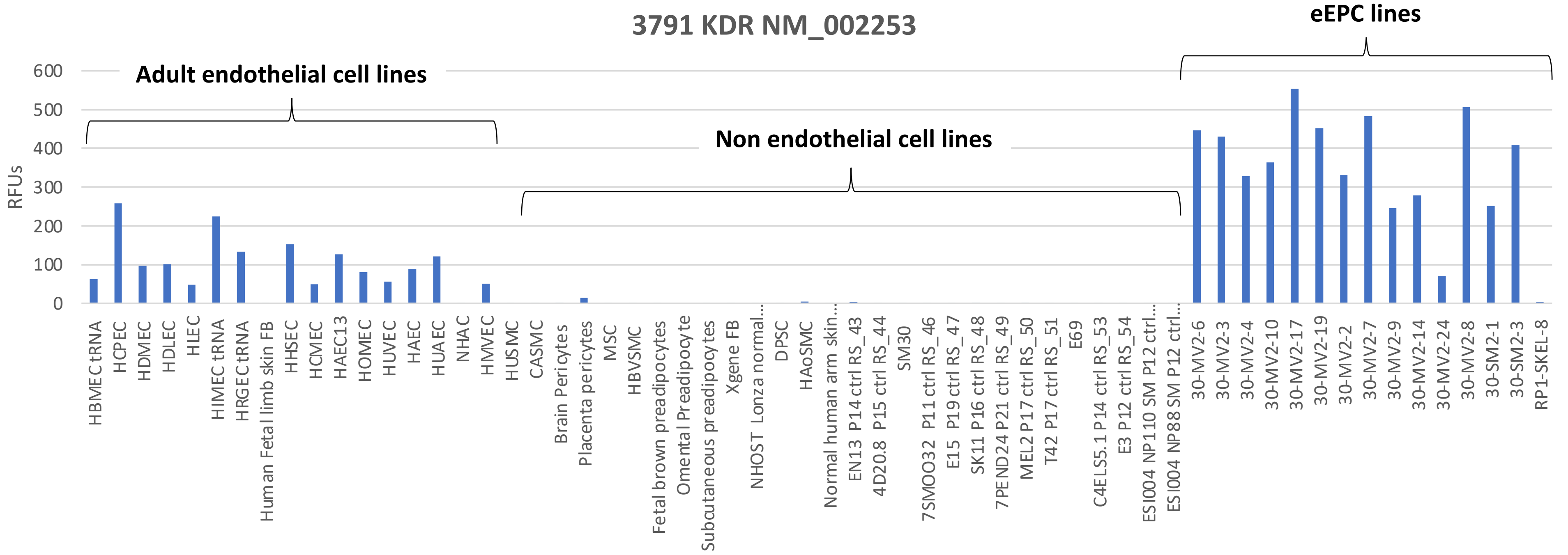


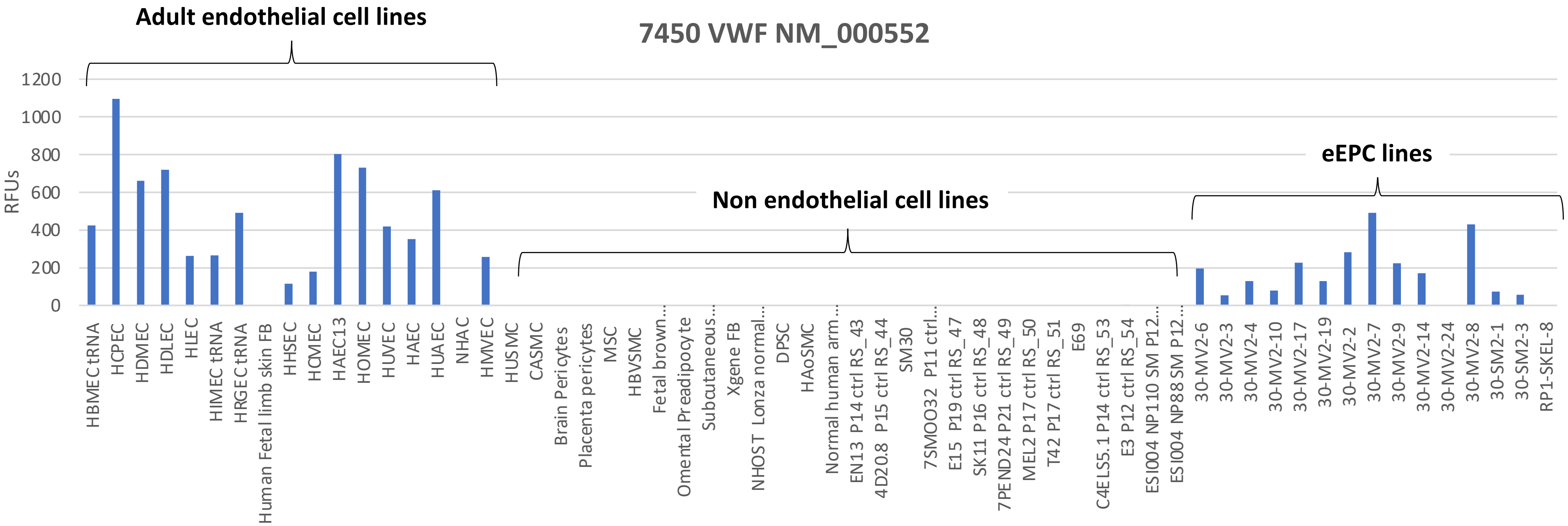

### Supplementary Figure 3. Endothelial specific gene expression in eEPC lines and adult EC lines (Related to Figure 3)

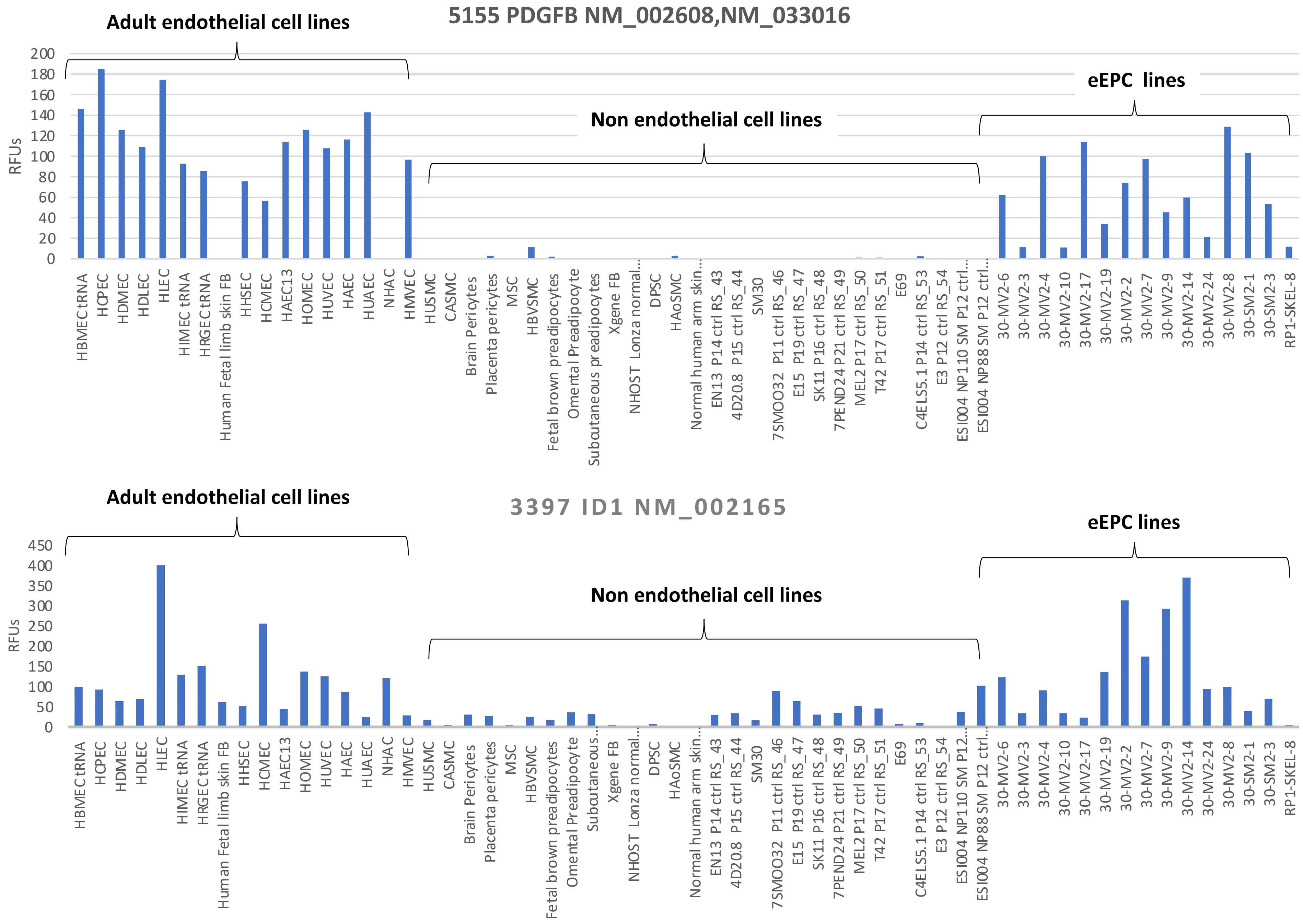

Supple Video 1A. 30MV2-4-RFP tube formation at 20% $O_2$

Control

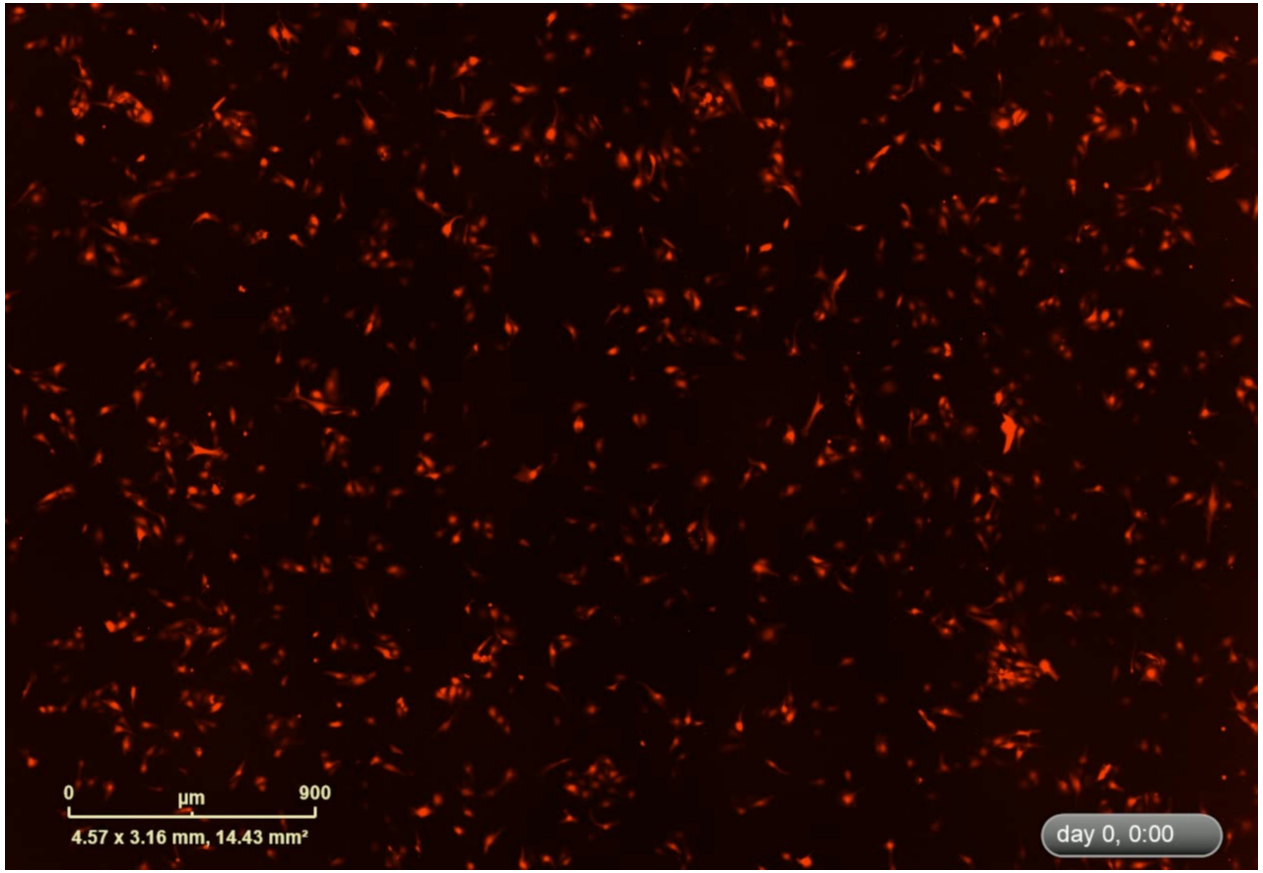


Control+VEGF

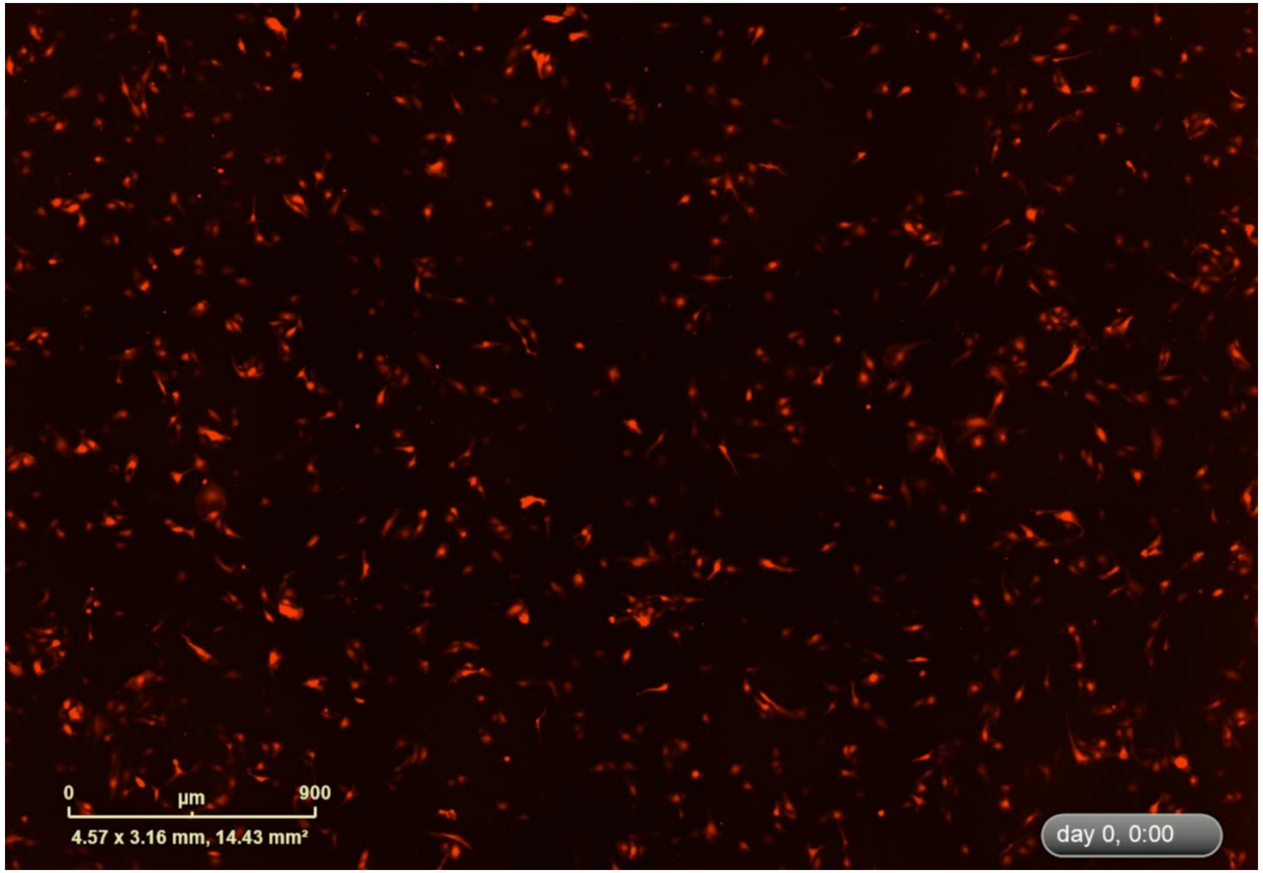

Supple Video 1B. 30MV2-14-RFP tube formation at 20% $O_2$

Control

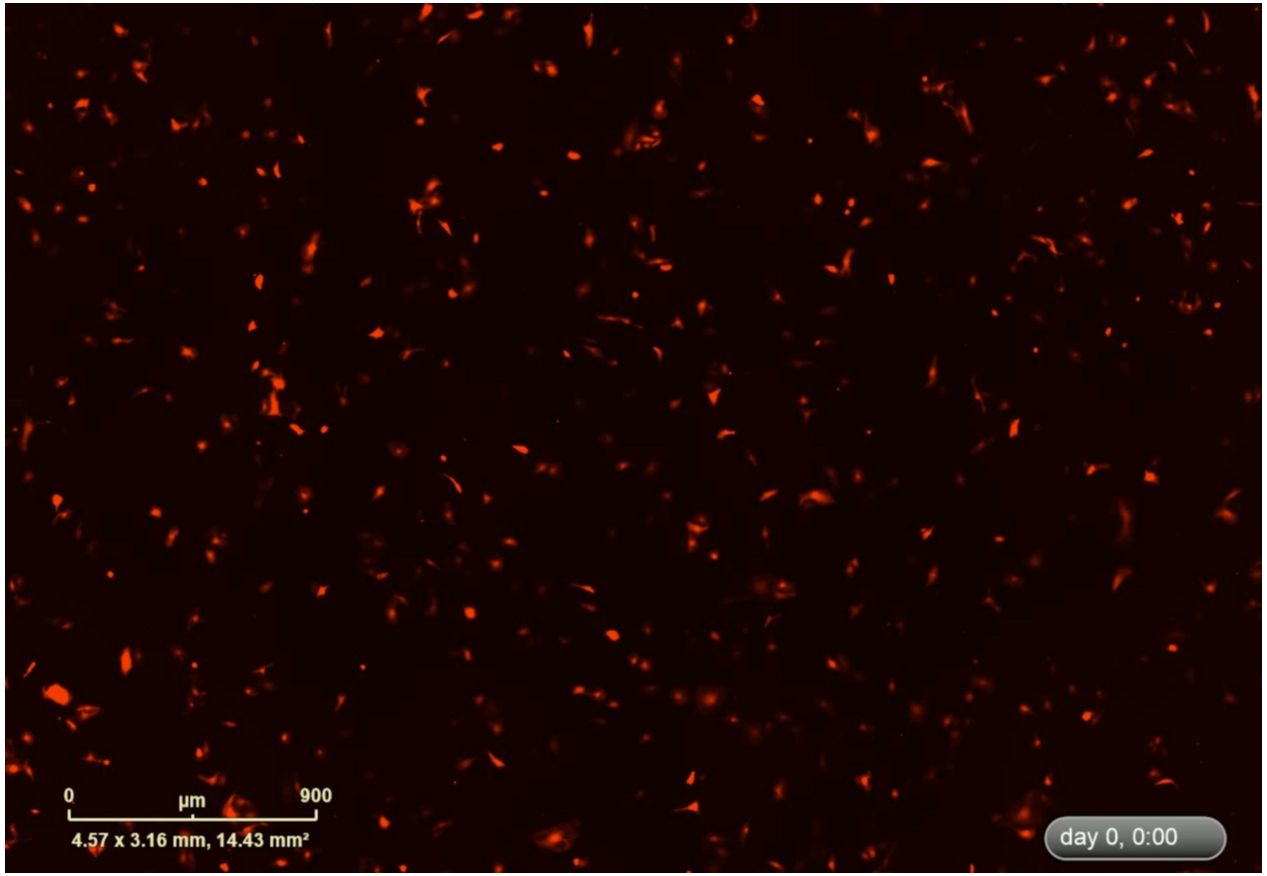


Control+VEGF

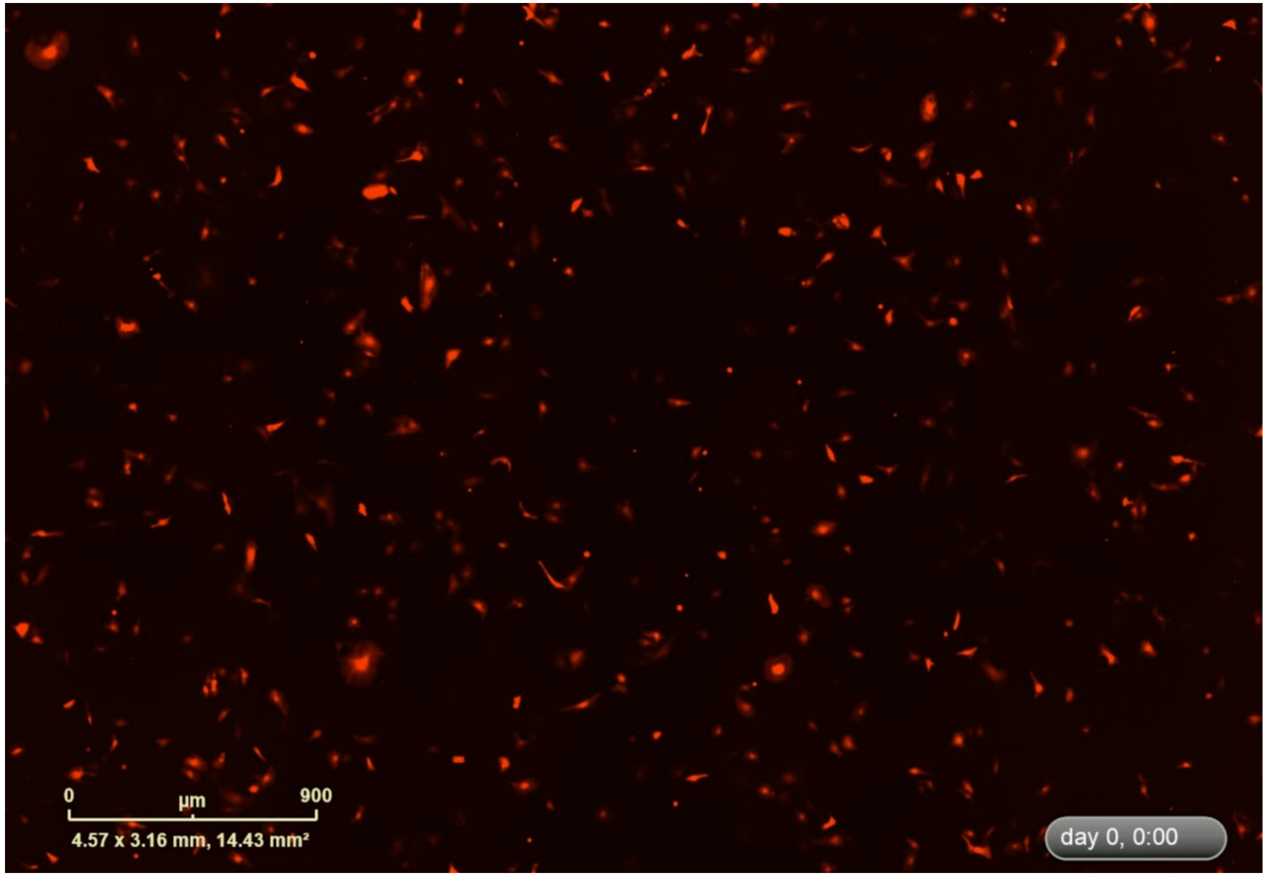

**Supplementary Figure 4.** Microarray analysis for global gene expression comparison between embryonic EC and adult EC lines, Filtered on Error- CV < 50.0 percent

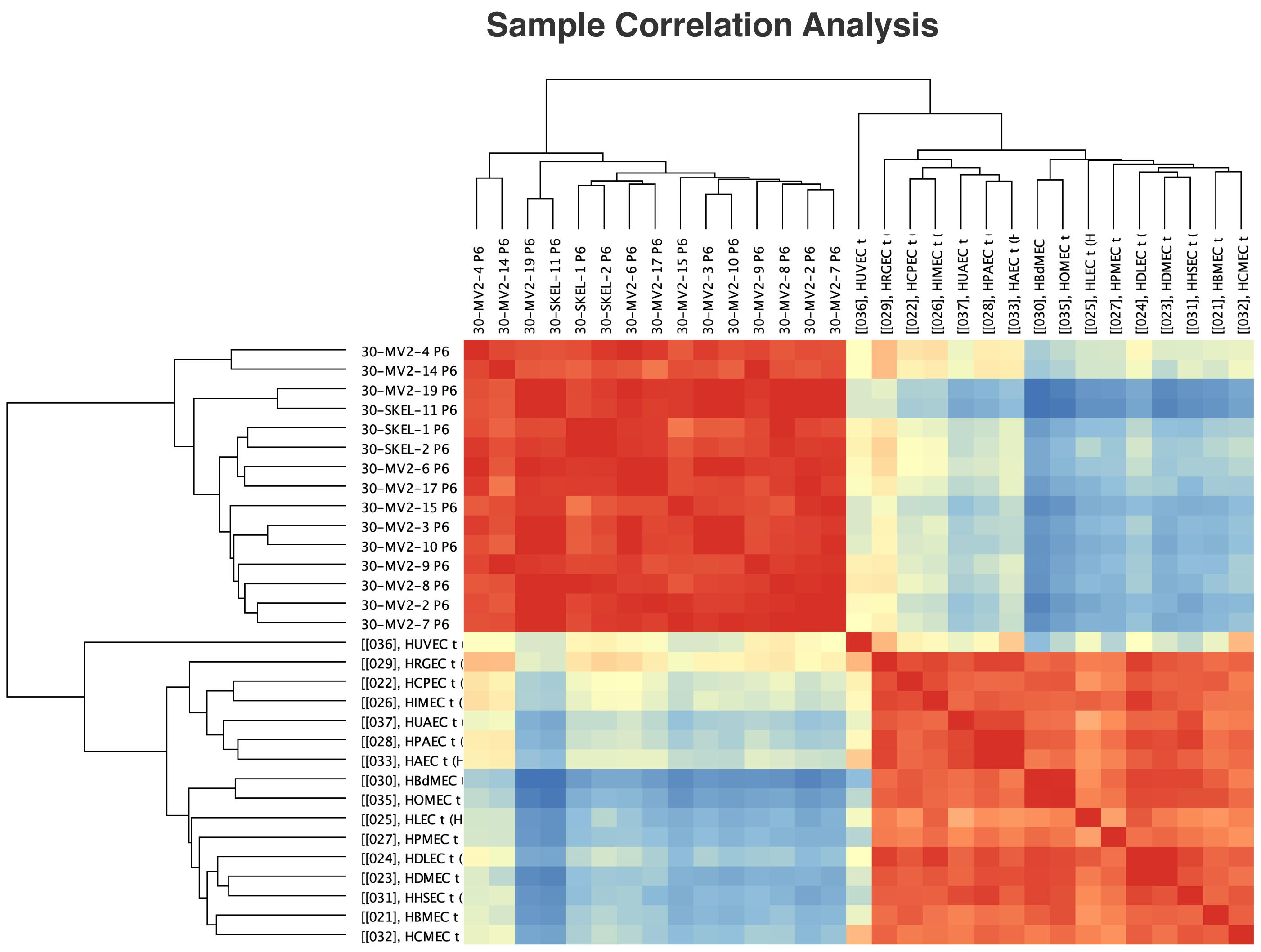


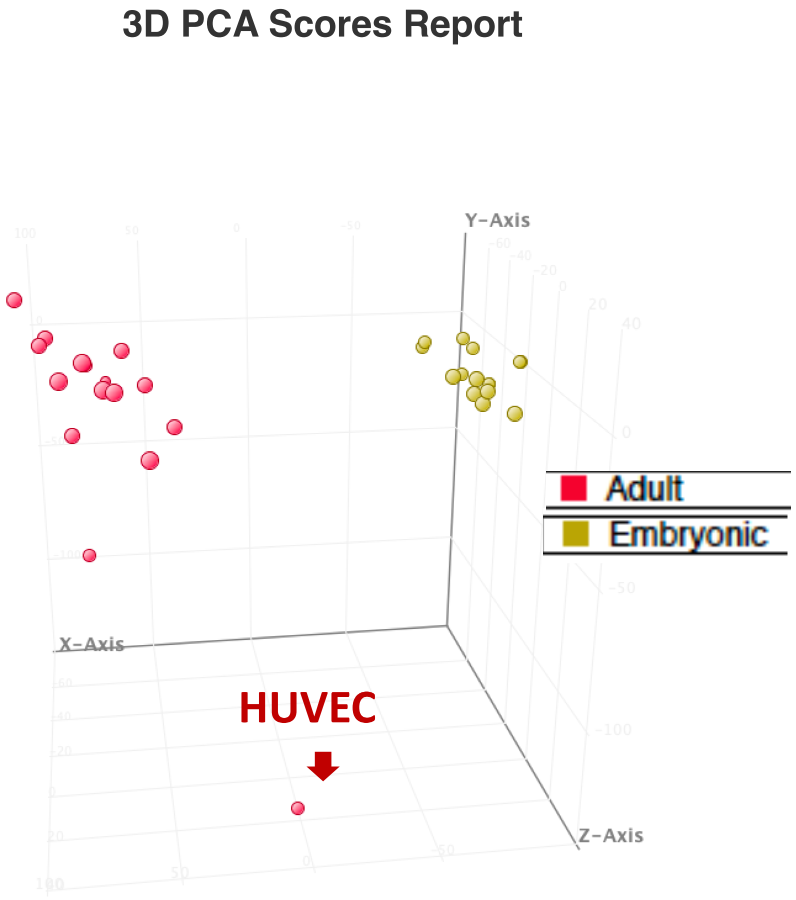

### Supplementary Figure 5A. Highly expressing genes expression in eEPC lines comparing to Adult EC lines (Related to Figure 7)

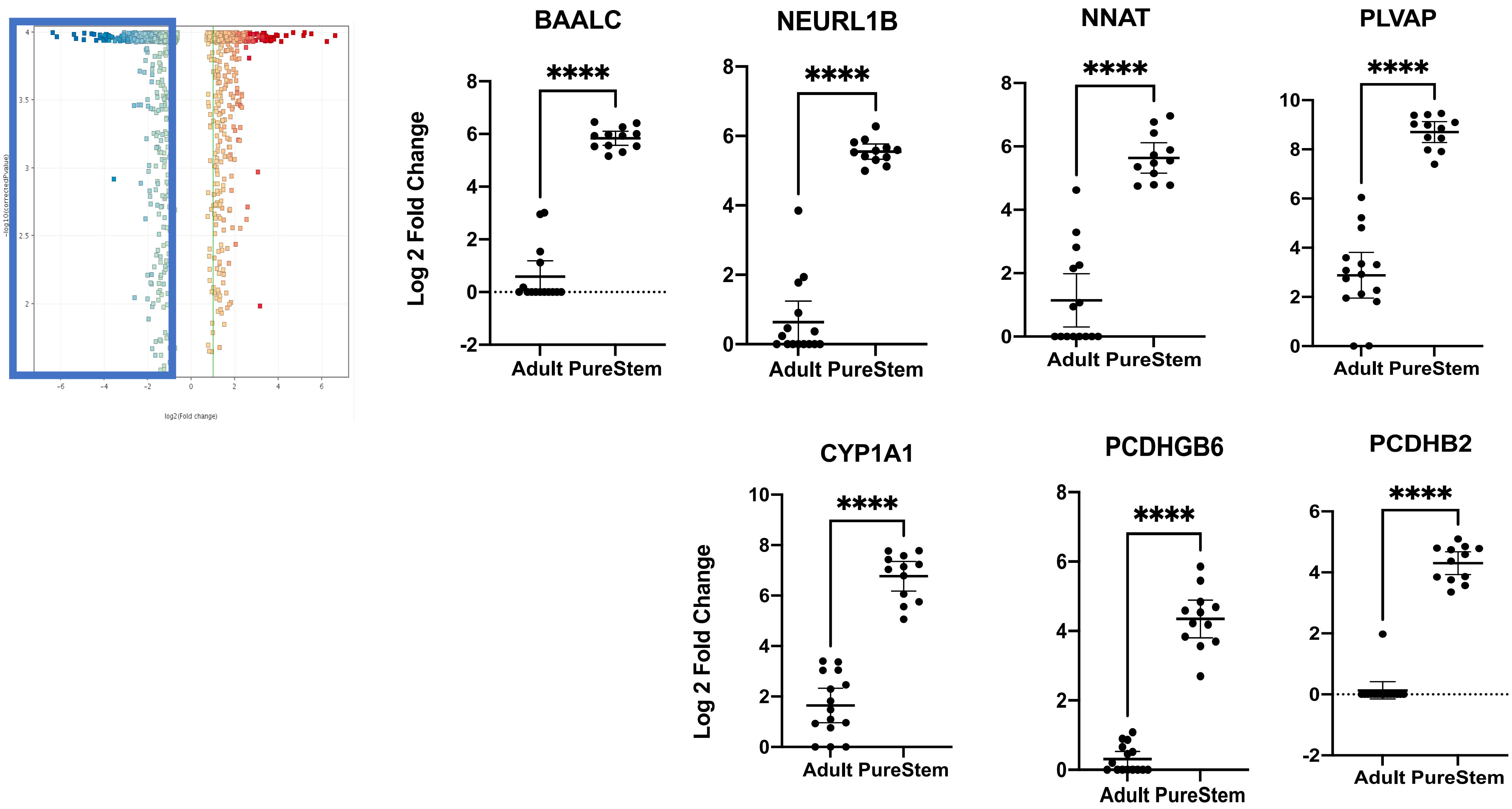

**Supplementary Figure 5B. Highly expressing genes expression in Adult EC lines comparing to eEPC lines (Related to Figure 7)**

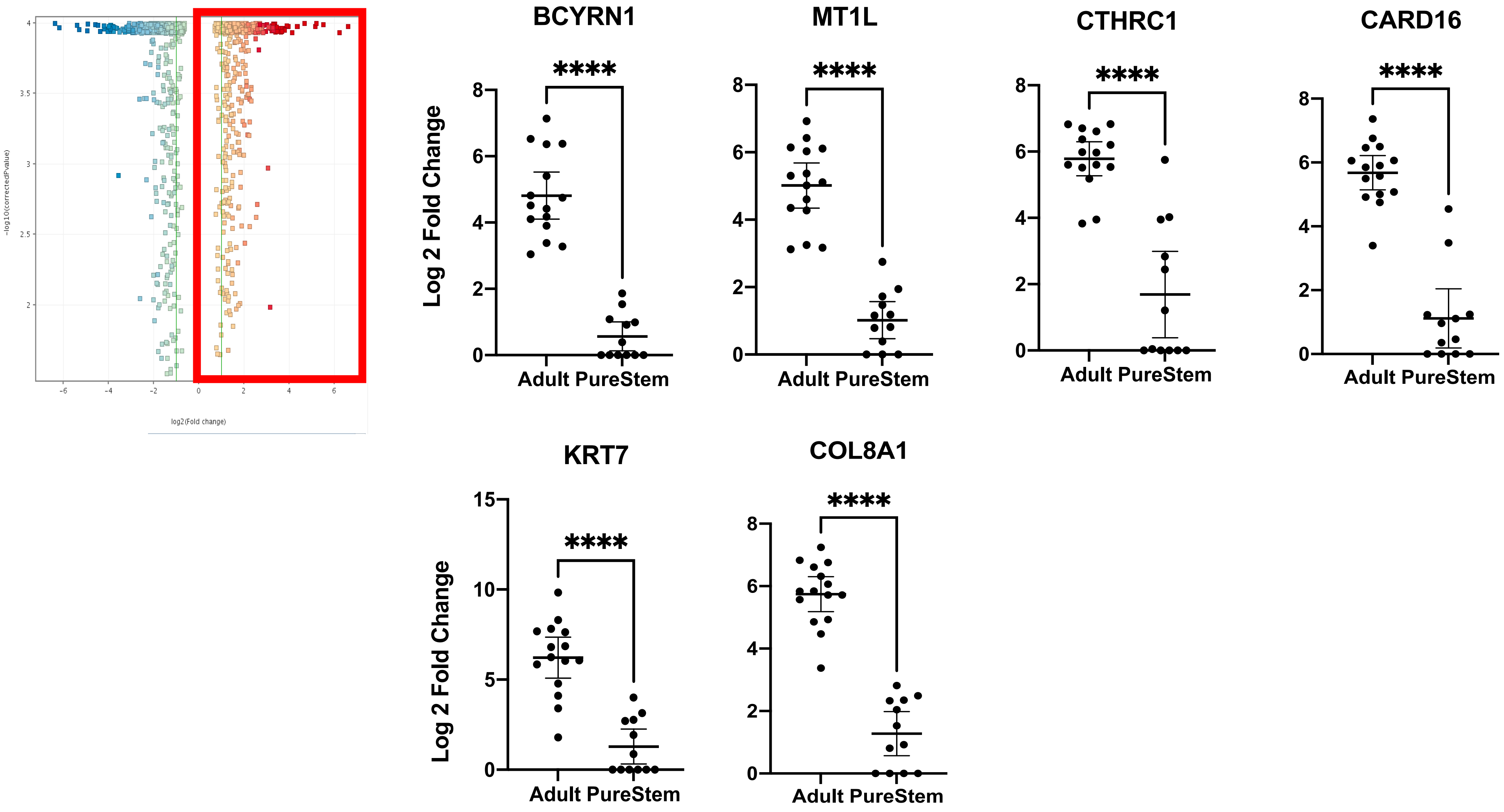

**Supplementary Figure 6.** eEPC exosomes have greater angiogenic potency than BM-MSC exosomes. **(A)** Angiogenic activity of eEPC line, 30-MV2-6 exosomes is higher than BM-MSC and PBS control ($p<0.01$; ANOVA)). **(B)** Dose response indicates 30-MV2-6 exosomes are 6-fold more potent than BM-MSCs. (n=3, biological replicate experiments in triplicate). **(C)** 50-fold enrichment of miR-126 in 30MV2-6 vs. BM-MSC by qPCR.

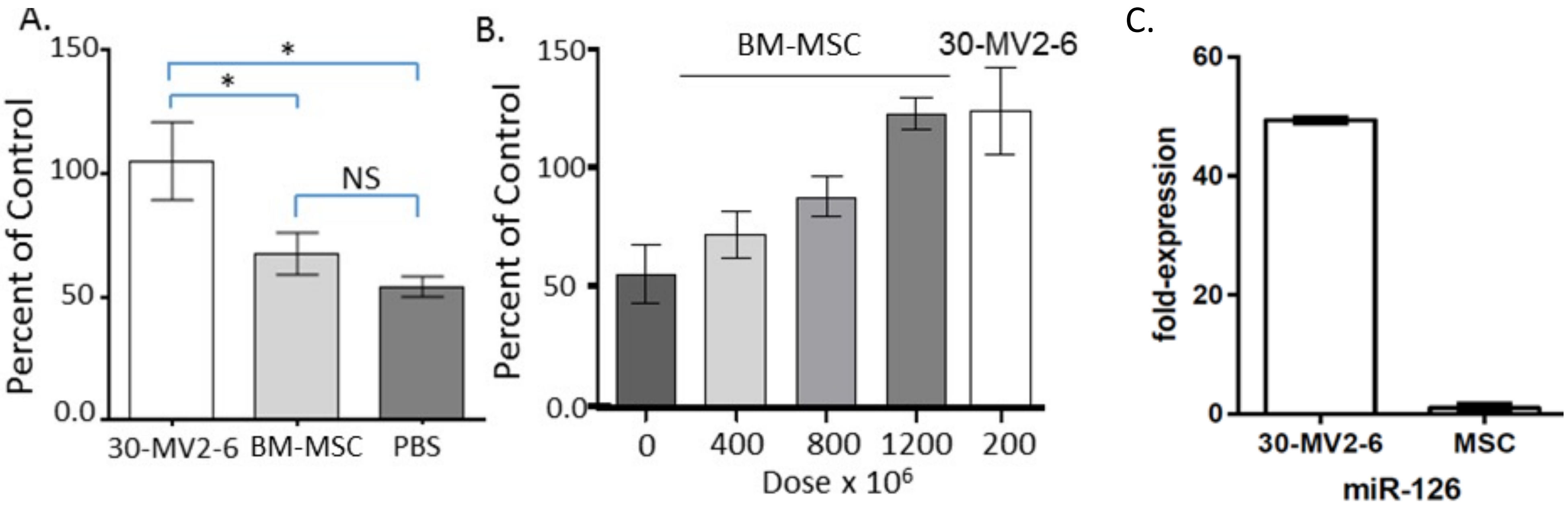

**Supplementary Figure 7.** Gene expression profile comparing HUVEC to 14 Adult Endothelial lines and 13 PureStem Endothelial lines

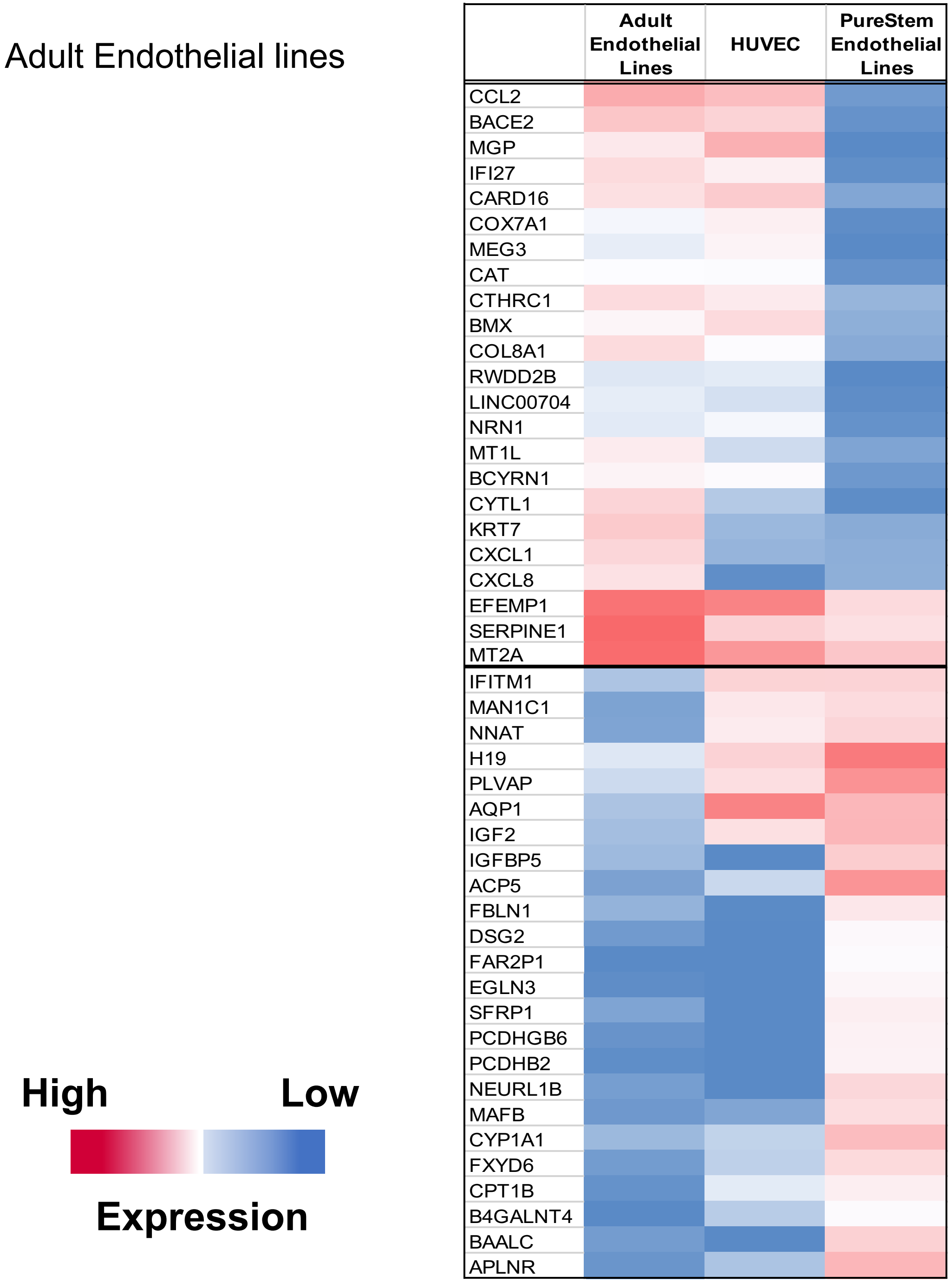
